# CRISPR-ChIP delineates a Menin-dependent oncogenic DOT1L complex in MLL- leukaemia

**DOI:** 10.1101/2023.03.17.533231

**Authors:** Omer Gilan, Charles C. Bell, Laure Talarmain, Daniel Neville, Kathy Knezevic, Daniel Ferguson, Marion Boudes, Yih-Chih Chan, Chen Davidovich, Enid Y.N. Lam, Mark A. Dawson

## Abstract

The regulation of all chromatin-templated processes involves the selective recruitment of chromatin factors to facilitate DNA repair, replication, and transcription. Chromatin immunoprecipitation (ChIP) is a critical experimental method used to provide spatiotemporal evidence for the coordination of these chromatin-based events including the dynamic regulation of chromatin modifications at cis-regulatory elements. However, obtaining a global appreciation of all the factors that influence a specific chromatin event has remained challenging. Here, as a proof of concept we demonstrate the utility of coupling unbiased functional genomics with ChIP to identify the factors associated with active transcription. Specifically, we use this method to identify the major chromatin factors associated with the catalysis of two evolutionarily conserved histone modifications; H3K4me3 present at the transcriptional start site and H3K79me2 present through the gene body of actively transcribed genes. With CRISPR-ChIP, we identify all the non-redundant COMPASS complex members required for H3K4me3 and demonstrate that RNA polymerase II is dispensable for the maintenance of H3K4me3. As H3K79me2 has a putative oncogenic function in leukaemia cells driven by MLL-translocations, using CRISPR-ChIP we reveal a functional partitioning of H3K79 methylation into two distinct regulatory units. An oncogenic DOT1L complex, where the malignant driver directs the catalytic activity of DOT1L at MLL-Fusion target genes and a separate endogenous DOT1L complex, where catalytic activity is directed by MLLT10. This functional demarcation provides an explanation for the observed synergy with Menin and DOT1L inhibitors and why Menin inhibition surprisingly controls methylation of H3K79 at a critical subset of genes that sustain MLL-fusion leukaemia. Overall, CRISPR-ChIP provides a powerful tool for the unbiased interrogation of the mechanisms underpinning chromatin regulation.

## Introduction

The transcriptional output of every gene is controlled by the cooperative function of chromatin modifiers, transcription factors and transcriptional co-activators. Our ability to unravel this complexity has emerged through a combination of biochemical approaches integrated with genomic assays such as ChIP-seq and RNA-seq^1^. However, the genomic assays are limited by the fact that they provide information for a single perturbation at a time. Similarly, as chromatin regulators often function within multi-subunit complexes, the biochemical approaches in isolation, are unable to delineate the precise complex components required for a chromatin-based event, such as a chromatin modification. The opportunity to simultaneously assess the contribution of thousands of proteins in the regulation of a defined chromatin-based event in an unbiased high-throughput manner may provide unique insights and accelerate our understanding of chromatin regulation.

To address this need we developed a ChIP-based CRISPR screen (CRISPR-ChIP) which enables the concurrent assessment of thousands of nuclear proteins in the coordinated regulation of chromatin. As a proof of principle, we chose to study the regulation of two histone modifications associated with active transcription. H3K4me3 is catalysed by the COMPASS family of methyltransferases and is primarily located over the transcriptional start site (TSS) of actively transcribed genes. In contrast, H3K79me2 is deposited by DOT1L in a non-redundant manner and is predominantly localised throughout the body of the transcribed locus. Notably, H3K79me2 is one of the few histone modifications directly implicated in cancer, specifically MLL-Fusion leukaemias, where several targeted therapies have been specifically developed and clinically translated to inhibit DOT1L and reduce H3K79methylation. Therefore, we sought to focus on this histone modification and study its regulation in MLL-Fusion leukaemia cells.

Using CRISPR-ChIP, which can couple regulation of chromatin events with a concomitant readout of cell viability, we reveal an unexpected discordance between the functional regulators of H3K79methylation and their essentiality in MLL leukaemia cells. Mechanistically, we demonstrate that underpinning this observation is a functional separation of DOT1L into two complexes in MLL leukaemia cells (i) an endogenous complex containing MLLT10 which deposits H3K79me2 at the majority of expressed genes and (ii) a neomorphic malignant DOT1L complex driven by the MLL-fusion oncogene which localises DOT1L to potentiate gene expression at a subset of target genes required for the maintenance of leukaemia. Additionally, our data provide detailed molecular insights into how Menin inhibitors unexpectedly and selectively influence H3K79methylation at a subset of genes and specify the mechanistic rationale for the synergistic efficacy of Menin and DOT1L inhibition in this disease.

## Results

### Establishing the CRISPR-ChIP method

Functional genomics using CRISPR screens have provided an unprecedented opportunity to uncover new biological insights in an unbiased manner. Thus far, the majority of CRISPR screening approaches have been restricted to positive and negative selection screens using cell viability as a read out^2^. More recently, these have expanded to FACS-based screens, which enable the selective enrichment of cells expressing a fluorescent protein or cell-surface target that is linked to a specific biological process^2^. As chromatin-based events cannot be easily assessed in this manner we sought to develop CRISPR-ChIP, which directly couples unbiased functional genomics with chromatin immunoprecipitation.

ChIP involves fragmentation of linker-DNA, which separates neighbouring nucleosomes, using sonication or enzymatic digestion^3^. Whilst micrococcal nuclease can provide mono- nucleosome resolution, methods that involve sonication of crosslinked chromatin usually result in di- and tri-nucleosome fragments^3^. Therefore, a key consideration in developing the method was the need to ensure that the integrated guide RNA (sgRNA) sequence remained coupled to the chromatin template that is enriched in a ChIP assay. To address this technical requirement, we chose to link the sgRNA 50-100bp upstream of the regulatory element being assessed in the ChIP assay (**Fig. 1a**).

**Figure 1:**
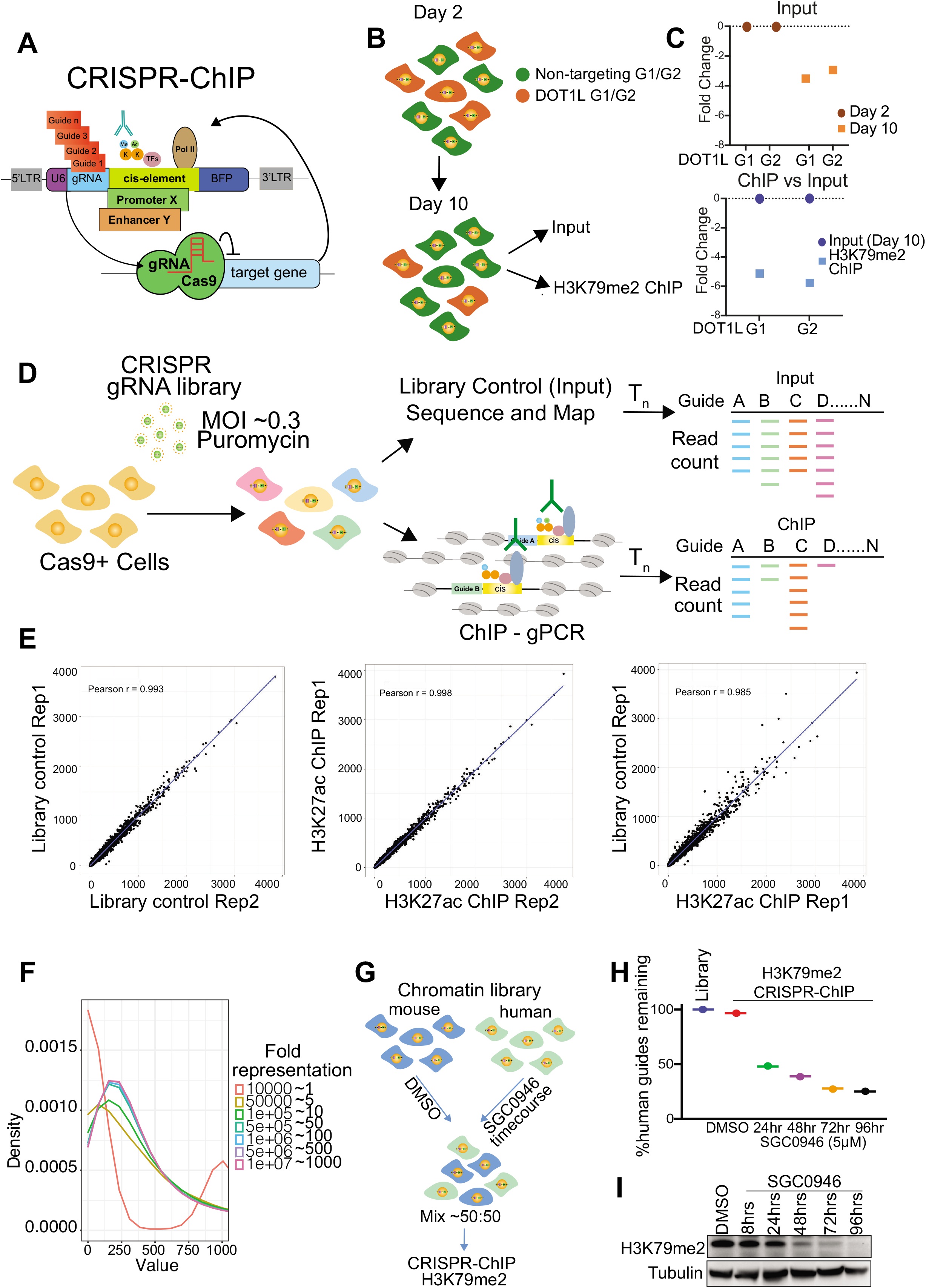
Establishing the CRISPR-ChIP method. **A)** Schematic of vector utilised for the screen highlighting the proximity of the gRNA sequence to the cis-regulatory element of interest driving expression of a fluorescent reporter. **B)** Schematic of CRISPR-ChIP proof-of-concept experiment with a mix of two independent DOT1L and two non-silencing guides sampled at day 2 and day 10 followed by ChIP-PCR NGS at day 10 for H3K79me2. **C)** H3K79me2 CRISPR-ChIP performed in murine MLL-AF9 cells at day 10 post-infection with a pool of two independent DOT1L and two independent control (non-silencing) guides. Fold-change depletion of DOT1L guides normalised to control guides in the library control sample (input) at day 2 and day 10 (Top panel). Fold-change depletion of DOT1L sgRNAs in the H3K79me2 ChIP at day 10 vs day 10 library control (input) (Bottom panel). Guide representation was assessed by next-generation sequencing (NGS). **D)** Schematic of the CRISPR-ChIP workflow. Cells are infected with a lentiviral CRISPR library (MOI <0.3) and cultured for a desired time period, cells were then collected as a library control or crosslinked for ChIP with antibody of interest. sgRNAs are amplified by PCR from ChIP DNA or library control (input) DNA and sequenced by NGS. Differential enrichment of sgRNAs between ChIP and library control is determined using MAGECK ^79^. **E)** The CRISPR- ChIP chromatin library was transduced into Cas9 negative K562 cells. H3K27ac ChIP was performed, and guide representation was assessed by NGS in two sampled replicates of library control and two independent ChIP samples. Data shown are correlation plots of library control Rep1 vs library control Rep2 (R=0.998), H3K27ac (50x10^6) Rep1 vs H3K27ac (50x10^6) Rep2 (R=0.993), and Library control Rep1 vs H3K27ac Rep1 (50x10^6) (R=0.985). **F)** Guide library was down sampled *in silico* at different fold representations, 10^7^ = ∼1000-fold representation down to 10000 guides = ∼1fold representation. Guide distribution was plotted for each sampling event, width of the distribution represents the degree of skewing, with a wider distribution being more skewed. **G)** Schematic of H3K79me2 CRISPR-ChIP of SGC0946 timecourse treatment in K562 cells infected with a human guide library and mixed with DMSO treated K562 cells infected with mouse guide library ∼ 1:1 ratio. **H)** Guide enrichment assessed by NGS and the percentage of human guides remaining in library control, H3K79me2 in the DMSO or SGC0946 treated samples at the indicated time points is shown. **I)** Western blot analysis of H3K79me2 levels after treatment with SGC0946 at the indicated timepoints, antibody against tubulin was used as a loading control.

To optimise our method, it was important to determine whether the lentivirally integrated regulatory sequences were modified and regulated in a similar manner to the corresponding endogenous loci. Following separate lentiviral infection, we sorted BFP+ cells expressing integrated regulatory sequences representing (i) a strong promoter (*EF1a* promoter), (ii) a weak promoter (*PGK* promoter), (iii) a developmental promoter (*Meis1* promoter) and (iv) the *Myc* enhancer^4^. We then performed ChIP-qPCR, to measure the enrichment of H3K27ac, H3K4me3, H3K9ac, H3K79me2, and Pol II at both the integrated regulatory sequence and the corresponding endogenous locus in human cells. Overall, our results showed that the chromatinised environment surrounding the virally integrated regulatory element undergoes a similar level of histone modification compared to the endogenous locus (**Extended Data Fig. 1a-d**). Next, we wanted to address if the histone modifications at our integrated reporter were regulated in a similar manner to the endogenous loci. As we were interested in assessing histone modifications associated with active transcription, we chose to use the full (∼1kb) EF1a promoter and assess H3K27ac and H3K79me2 as the enzymes that deposit these modifications are amenable to inhibition with potent and selective small molecules^5, 6^. This allowed us to assess the changes in the level of these histone modifications around our integrated promoter sequence and the corresponding endogenous locus. We treated cells expressing the integrated EF1a promoter sequence with a DOT1L inhibitor (SGC0946) or a P300/CBP inhibitor (A-485) and repeated our ChIP-qPCR analysis. Our data confirmed that treatment of the cells with SGC0946 and A-485 resulted in a marked decrease in H3K79me2 and H3K27ac respectively, at both the integrated promoter sequence and at endogenous genes marked by the histone modifications (**Extended Data Fig. 1e-f**). Together these results suggested that our integrated promoter sequence was subjected to the same epigenetic regulatory processes active at endogenously expressed genes.

Having established the suitability of the integrated promoter sequence to assess changes in histone modifications associated with gene expression we next sought to validate the ability to discern the depletion of sgRNAs against nuclear proteins that regulate a histone modification of interest. Cognisant of the fact that many chromatin regulators are also common-essential proteins, we wanted to understand if we were able to capture major regulators of a histone modification that were also required for the viability of the cell. To achieve this, we mixed two DOT1L guides with two non-targeting guides and introduced these into leukaemia cells sustained by MLL-AF9 (**Fig. 1b**). We then harvested the cells at both day 2 and day 10 after sgRNA infection. Since H3K79me2 has no known demethylase and we had previously shown that this modification is slowly turned over^7^, we waited until day 10 to perform ChIP for H3K79me2 to allow for marked decrease in H3K79me2 (**Fig. 1b**). To assess the effects on cell viability, we determined the relative guide abundance of the guides at day 10 relative to day 2. As expected, we found that DOT1L guides were depleted in MLL-Fusion leukaemia cells at day 10 (**Fig. 1c and Extended Data Fig. 1g**). To determine the impact on H3K79me2, we performed a ChIP assay at day 10 in viable cells and compared the representation of the sgRNAs from the ChIP sample relative to the input sample at day 10. Here, we see a marked depletion in DOT1L sgRNAs from the H3K79me2 ChIP material, suggesting that in principle, the CRISPR-ChIP method can reliably identify a known epigenetic regulator of a histone modification even when the regulator is required for cell viability (**Fig. 1c and Extended Data Fig. 1g**).

Another important consideration in developing the method to be suitable for a high-throughput screen with a library of thousands of sgRNAs is that the absolute amount of DNA that is immunoprecipitated in a typical ChIP assay can be relatively low (∼0.1-10% of the input DNA for the locus of interest). Therefore, unlike conventional CRISPR screening approaches which harvest the entire cellular genomic DNA, the lower yield in this method is a critical parameter that could directly influence the representation of guides in the library. To establish the technical feasibility of the method, we developed a guide RNA library targeting 1144 chromatin and transcriptional regulators (each gene was targeted by 6 guides each) (**Supplementary Table 1**). In our workflow (**Fig. 1d**), the lentiviral sgRNA library is infected into cells expressing Cas9; importantly, we aim for a low multiplicity of infection (MOI ∼0.3) to ensure that only a single sgRNA is integrated per cell. Therefore, we are only assessing the contribution of genetic depletion of a single chromatin regulator per cell. Expression of BFP from the integrated reporter enables FACS sorting of cells expressing the reporter construct and sgRNA. The viable BFP+ cells are then harvested at the desired timepoint and from the same population of cells, we apportion one sample to provide us with the input library, while the remaining cells are cross-linked, sonicated and taken forward in a standard ChIP assay for the target of interest (**Fig. 1d**). Both input and ChIP samples are then assessed by next-generation sequencing (NGS) for the depletion or enrichment of guides (using MAGECK) in the ChIP sample relative to input.

To assess the cell number required to reliably retain library representation, we performed CRISPR-ChIP for H3K27ac from Cas9 negative K562 cells. Since no gene editing occurs, we could determine the extent to which library representation is altered in the ChIP sample (relative to the input library) from different starting cell numbers. Our results revealed that library representation following lentiviral infection was very highly correlated between two independent replicates of the input library (**Fig. 1e**). Importantly, we also found that the two independent replicates of H3K27ac ChIP were highly correlated (**Fig. 1e**). Notably, an excellent correlation in the representation of the sgRNA library between the ChIP and input samples was faithfully maintained from 100 million cells (r=0.985) down to 5 million cells (r=0.958) (**Fig. 1e and Extended Data Fig. 2a**). As ChIP efficiency is dependent on the quality of the antibody and abundance of the target of interest, we wanted to use our data to model the point at which technical limitations in the ChIP could result in substantial skewing in library representation. To address this, we performed *in silico* down sampling of the H3K27ac ChIP sample and plotted the guide representation density. This analysis revealed that substantial skewing only begins to occur when the ChIP sample contains less than 50-fold representation of each sgRNA within the library (**Fig. 1f**).

**Figure 2:**
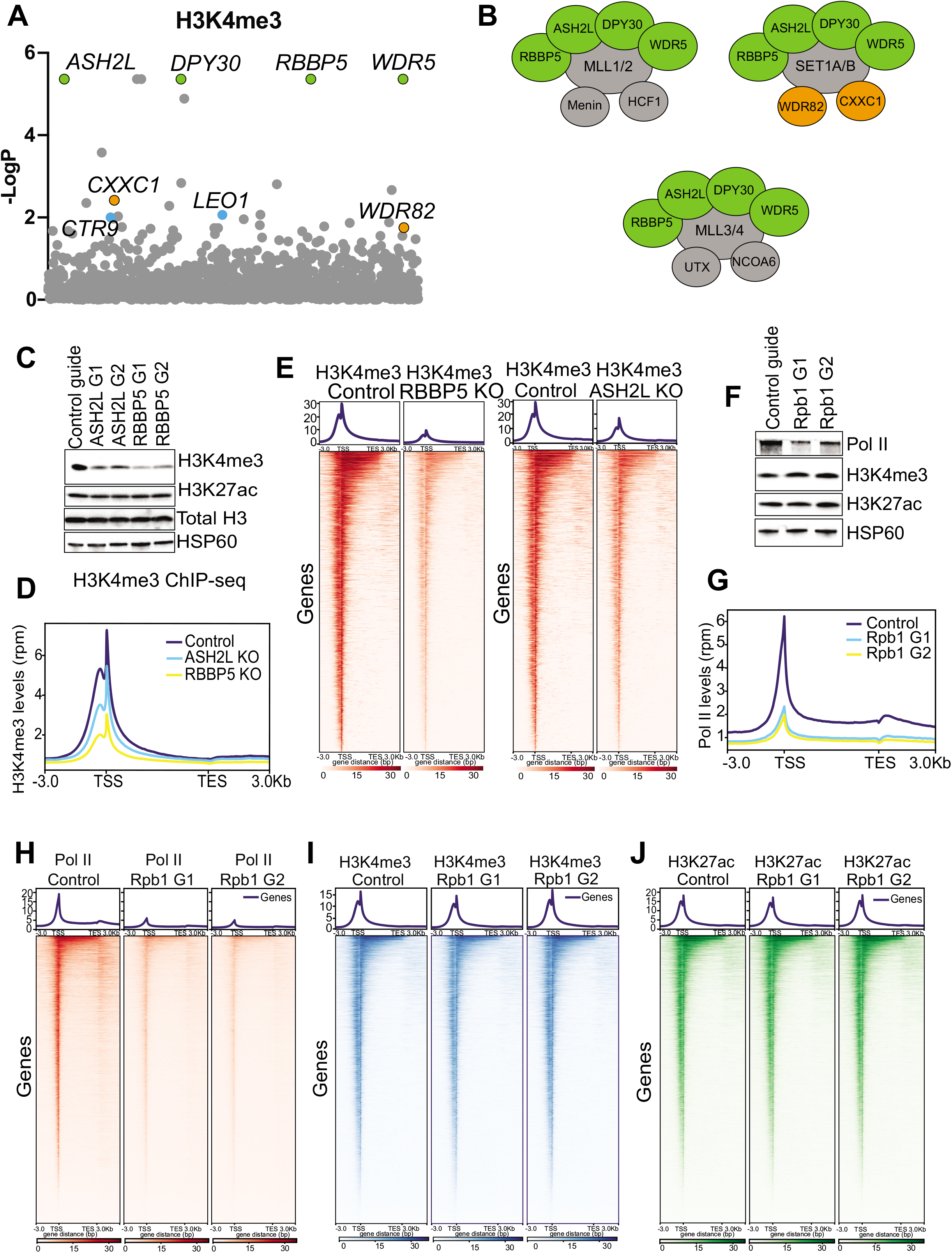
Characterising the regulation of H3K4me3 using CRISPR-ChIP. **A)** CRISPR-ChIP bubble plot showing the genes required for the maintenance of H3K4me3 at the EF1a promoter in K562 cells. P-values calculated using MAGECK algorithm. **B)** Schematic of the COMPASS and COMPASS-like complexes highlighting subunits identified in the CRISPR-ChIP screen, WRAD complex in green and SET1A/B complex specific subunits in orange. **C)** Western blot analysis of K562 Cas9 cells infected with control guide (Safe) or two independent guides targeting ASH2L or RBBP5 using antibodies against H3K4me3, H3K27ac, Total H3 and HSP60 (loading control). **D)** Profile plot of H3K4me3 ChIP-seq in K562 Cas9 cells transduced with control (NT), ASH2L sgRNA or RBBP5 sgRNA. **E)** ChIP-seq heatmap of H3K4me3 in K562 Cas9 cells infected with control guides or guides targeting RBBP5, ASH2L. Showing all genes and ranked based on H3K4me3 levels in control cells. **F)** Western blot analysis of K562 Cas9 cells infected with control guide (Safe) or two independent guides targeting Rpb1 (largest subunit of RNA Pol II) using antibodies against Rpb1, H3K4me3, H3K27ac and HSP60 (loading control). **G)** Profile plot of Pol II ChIP-seq in K562 Cas9 cells transduced with control (NT), Rpb1 sgRNA (G1) or Rpb1 sgRNA (G2). **H)** ChIP-seq heatmap of RNA-Pol II in K562 Cas9 cells infected with control guides or two independent guides targeting Rpb1. Showing all genes and ranked based on RNA-Pol II levels in control cells. **I)** ChIP-seq heatmap of H3K4me3 in K562 Cas9 cells infected with control guides or two independent guides targeting Rpb1. Showing all genes and ranked based on H3K4me3 levels in control cells. **J)** ChIP-seq heatmap of H3K27ac in K562 Cas9 cells infected with control guides or two independent guides targeting Rpb1. Showing all genes and ranked based on H3K4me3 levels in control cells.

The change in abundance of histone modifications is not binary and instead spans a continuous spectrum, we therefore wanted to understand whether CRISPR-ChIP has sufficient sensitivity to capture a quantitative change in the levels of the modification. To achieve this, we infected separate Cas9 negative K562 cells with either a human or mouse guide library and treated the cells harbouring the human library with a DOT1L inhibitor (SGC0946) for up to 96hrs. In parallel, we treated the cells containing the mouse library with vehicle (DMSO). The mouse and human K562 cells were then mixed at 1:1 ratio and the relative guide enrichment from the ChIP for H3K79me2 were determined at various timepoints. This approach allowed us to compare the enrichment of the CRISPR library from SGC0946 treated cells where there is a gradual and progressive reduction in H3K79me2 (**Fig. 1g**) compared to vehicle treated cells where the modification stays constant. Our results showed that the CRISPR-ChIP for H3K79me2 in the treated/vehicle (human/mouse) mix accurately reflected the input guide distribution and enrichment (**Fig. 1h**). Furthermore, we clearly observed a quantitative time-dependent reduction in the representation of the human library with our ChIP for H3K79me2 following SGC0946 treatment relative to the vehicle treated mouse library (**Fig. 1h**). Notably, these data closely parallel the change in abundance noted by western blot (**Fig. 1i**). Together these data serve to highlight the fact that CRISPR-ChIP can detect a quantitative alteration in the level of a chromatin modification. Finally, we also performed MAGECK analysis on the CRISPR-ChIP data from Cas9 negative cells to determine the expected background scores generated by the method when no editing occurs (**Extended Data Fig. 2b**). This control analysis revealed the low probability of false positives arising from CRISPR-ChIP providing further confidence that our method was robust.

### CRISPR-ChIP identifies the major non-redundant regulators of H3K4me3

The tri-methylation of lysine 4 on histone H3 (H3K4me3) is one of the best characterised histone modifications, as it is invariably localised at the transcription start site of actively transcribed genes in all eukaryotes and its abundance is directly correlated with transcriptional activity at the locus^8, 9^. The yeast model organism *Saccharomyces cerevisiae* has a single methyltransferase, SET1, that deposits this modification. However, in mammals there are six H3K4 methyltransferases (SET1A/B and MLL1-4), which have all been shown to strongly associate with the initiated form of RNA polymerase II (Pol II)^10, 11^. Of these, SET1A/B are thought to be the methyltransferases responsible for the majority of H3K4me3^12^, whereas, MLL1/2 deposit H3K4me3 at key developmental genes^13^. The methyltransferases in isolation are not efficient enzymes at depositing H3K4me3, instead their catalytic activity is almost entirely dependent on their incorporation into a multi-subunit protein complex called COMPASS (containing either SET1A/B) or COMPASS-like (containing one of MLL1-4). The various subunits of the COMPASS and COMPASS-like complexes are essential for the stability and functional integrity of the methyltransferases.

To uncover the non-redundant functional regulators of H3K4me3, the prototypical histone modification associated with active transcription, we performed CRISPR-ChIP for H3K4me3 in K562 cells (**Fig. 2a and Supplementary Table 2**). The top hits in our screen were the four proteins commonly referred to as WRAD (**Fig. 2b**), including WD repeat-containing protein 5 (WDR5), Retinoblastoma Binding Protein 5 (RBBP5), Absent-Small-Homeiotic-2-Like protein (ASH2L) and Dumpy-30 (DPY30)^14^. The four WRAD components are fundamental core proteins that comprise all COMPASS and COMPASS-like complexes and are known to be required for the functional integrity of COMPASS and COMPASS-like complexes, which deposit and maintain of H3K4me3^13, 14^. In line with the results of the CRISPR-ChIP screen, we subsequently validated that CRISPR-Cas9 mediated deletion of ASH2L or RBBP5 resulted in a global loss of H3K4me3 by western blot and ChIP-seq (**Fig. 2c-e and Extended Data Fig. 2c)**. Consistent with the fact that there is significant redundancy between the catalytic orthologs SETD1A/B, MLL1/2 and MLL3/4, none of these H3K4 methyltransferase enzymes were identified in our screen. However, the screen did identify CXXC1 and WDR82 that are specific subunits of the mammalian SETD1A/B COMPASS complex (**Fig. 2a-b**). These findings are entirely consistent with biochemical and cell biology studies showing that loss of CXXC1 and WDR82 is followed by a very specific global loss of H3K4me3 and not H3K4me1 or H3K4me2^15–17^. A further testament to the validity of this screening approach is the fact that we also identified several members of the mammalian PAF complex which has been carefully characterised through various biochemical approaches in *S. cerevisiae* and shown to interact with Pol II to provide a scaffold for the COMPASS complex and consequent deposition of H3K4me3^18^.

### Loss of RNA Pol II does not alter global levels of H3K4me3

The reciprocal interplay between Pol II and H3K4me3 has been the subject of intense debate^19–21^. It has been proposed that rather than being instructive for transcription, H3K4me3 may be a consequence of it, and instead, plays a key role in influencing transcriptional consistency^22^ and regulating co-transcriptional events such as splicing and termination of transcription^23, 24^. In this regard, a striking observation from the screen was the clear absence of Pol II subunits and the general transcriptional machinery (**Fig. 2a**). To assess whether these are false negative hits, we independently verified the finding that loss of Pol II did not affect global levels of H3K4me3 by depleting the major subunit of Pol II, Rpb1, using two independent sgRNAs. Western blot analysis revealed that depletion of Rpb1 did not impact the global levels of H3K4me3 (**Fig. 2f**). As a distinct control, we included H3K27ac, another dynamic and rapidly turned over histone modification associated with gene activity. Here again, we found that loss of Rpb1 did not affect global levels of H3K27ac (**Fig. 2f**). To confirm these results with an orthogonal approach, we performed ChIP-seq for both Pol II, H3K4me3 and H3K27ac after depletion of Rpb1. These data again confirm that H3K4me3 and Pol II levels at TSS are highly correlated in the control cells (**Extended Data Fig. 2d**). Furthermore, as expected, depletion of Pol II by CRISPR-mediated knockout of Rpb1 resulted in a profound reduction of Pol II at chromatin (**Fig. 2g-h**). However, again we found that the dramatic loss of chromatin bound Pol II had a negligible effect on H3K4me3 genome-wide (**Fig. 2i**).

It has been suggested by studies in yeast that H3K4me3 levels can persist for several hours after transcription ceases^25^. Therefore, to ascertain whether the retention in H3K4me3 levels after Pol II deletion is also seen with other histone modifications associated with active transcription, we also performed ChIP-seq for H3K27ac following Pol II depletion. Consistent with our observations for H3K4me3, Pol II was also not required for the genome-wide maintenance of H3K27ac (**Fig. 2j**). Taken together, these results suggest that although H3K4me3 is strongly associated with active transcription, a global decrease in chromatin bound Pol II has little influence on H3K4me3 levels.

### MLLT10 and global H3K79me2 are dispensable for survival of MLL-Fusion leukaemia cells

We were encouraged by the fact that in a single experiment, CRISPR-ChIP had identified most of the known regulators of H3K4me3. Therefore, we next wanted to apply the CRISPR-ChIP approach to study the regulation of a histone modification closely associated with the maintenance of malignant gene expression in a unique cancer setting, leukaemias driven by MLL-fusion oncogenes^26^. H3K79 lies in the globular domain of histone 3 and unlike H3K4me3, which is deposited by multiple enzymatic subunits, H3K79methylation is known to be globally regulated by a single enzyme, which in mammals is DOT1L^27^. H3K79me2 is distributed throughout the coding region of actively transcribed genes, where it is thought to play a role in facilitating transcription. Previous proteomic and biochemical studies have shown that the endogenous DOT1L complex is comprised of DOT1L, MLLT10, MLLT1/ENL and MLLT6/AF17^7, 28^ (**Extended Data Fig. 3a**). Moreover, whilst MLLT1/ENL is also found in the P-TEFb/super elongation complex (SEC), the other components of the DOT1L complex (DOT1L, MLLT10 and MLLT6) are mutually exclusive with P-TEFb/SEC^28–30^. On the other hand, in leukaemia cells containing an MLL-Fusion oncogene, DOT1L has been shown to be recruited by MLL-Fusion partners to increase H3K79methylation at critical MLL-Fusion target genes associated with leukaemia development^31–34^. Whilst DOT1L activity in MLL-Fusion leukaemia has been intensively studied^35^, the relative contribution of the MLL-Fusion oncogene to the regulation of H3K79me2 and whether there are other major regulators of this histone modification remains unclear.

**Figure 3:**
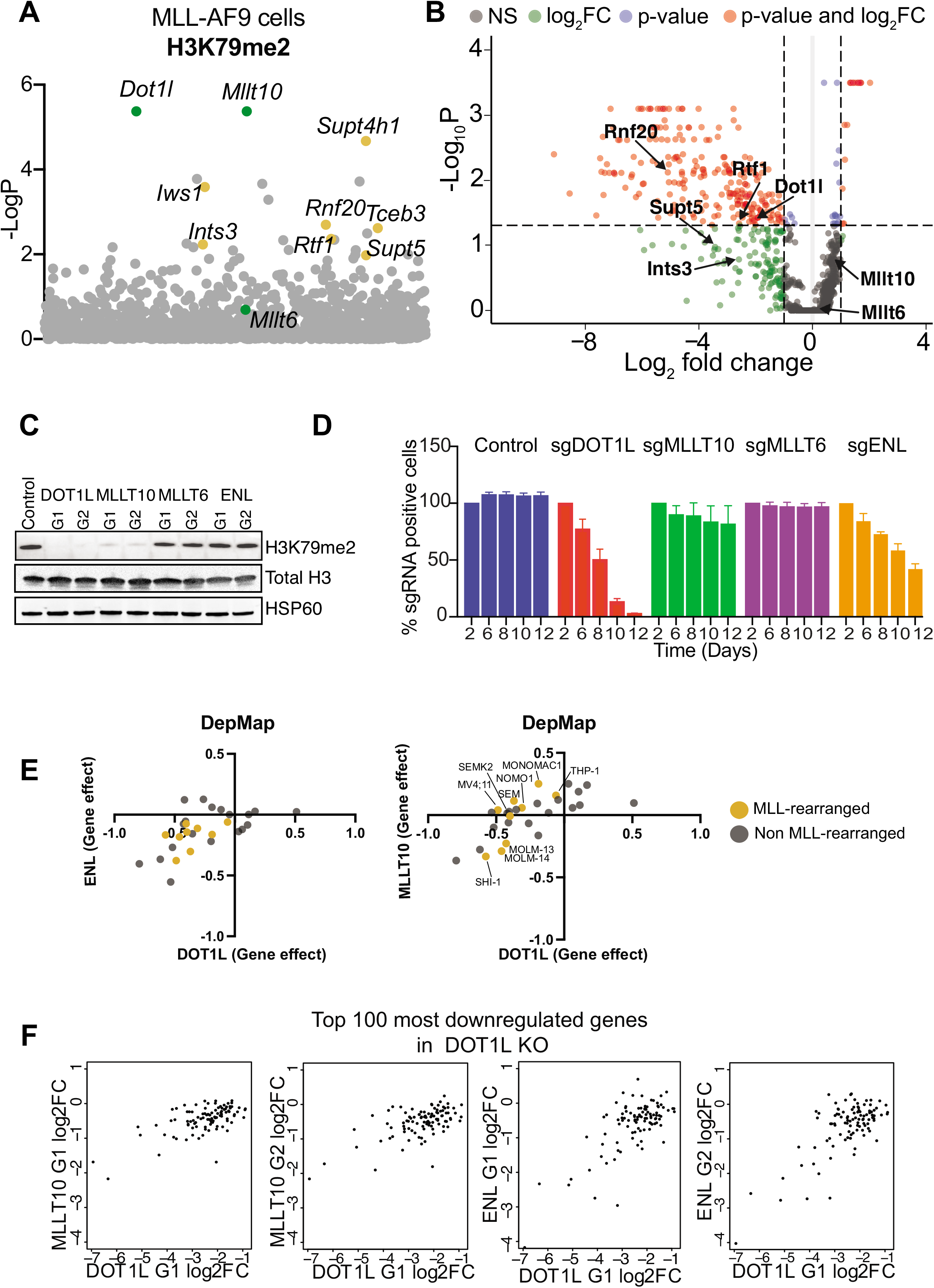
MLLT10 and global H3K79me2 are dispensable for the survival of MLL-Fusion leukaemia cells. **A)** Bubble plot showing the genes required for the maintenance of H3K79me2 at the EF1a promoter in the CRISPR-ChIP vector in murine MLL-AF9 cells. P-values calculated using MAGECK algorithm. **B)** Scatter plot of gene dependencies in murine MLL-AF9 cells as assessed from day 14 of the CRISPR-ChIP screen, highlighted members of the DOT1L complex (DOTCOM) contained in the library, DOT1L, MLLT10 and MLLT6. **C)** Immunoblot analysis of H3K79me2, Total H3 and HSP60 in MLL-AF9 Cas9 cells transduced with the indicated sgRNAs. **D)** sgRNA negative selection competition assays in MLL-AF9 Cas9 cells transduced with either a non-silencing guide (Control) or guides targeting DOT1L, MLLT10, MLLT6 or ENL. Percentage of sgRNA positive cells remaining over time. Data represents n=3 experiments. Mean +/- sd. **E)** Analysis of DepMap cell line dependencies comparing MLLT10 vs DOT1L gene effect and MLLT1(ENL) vs DOT1L gene effect across wild-type MLL and MLL-rearranged acute leukaemia cell lines. Cell lines containing MLL rearrangements are highlighted in orange. **F)** Correlation plot of RNA-seq LFC between the indicated knockouts (MLLT10 and ENL) for the top 100 downregulated genes vs DOT1L KO cells.

To address these questions, we performed a CRISPR-ChIP screen for H3K79me2 in MLL-AF9 leukaemia cells. Our results clearly identified DOT1L and MLLT10 but not the other DOT1L complex members as key non-redundant regulators of H3K79me2 (**Fig. 3a and Supplementary Table 3**). Early biochemical studies in yeast established the importance of H2Bub for the methylation of H3K79 by DOT1L^36–38^ and more recently the mammalian ring finger protein 20 (RNF20), the major H2B ubiquitin ligase, was shown to be required for the maintenance of H3K79me2 and the DOT1L mediated malignant gene expression program in MLL-AF9 leukaemia cells^39^. In line with these findings, CRISPR-ChIP also identified RNF20 as a major non-redundant regulator of H3K79me2 (**Fig. 3a**). Whilst the PAF complex is known to be required for efficient deposition of H3K79me2, our CRISPR-CHIP data specifically identified RTF1, which in contrast to its role in yeast, is a labile PAF complex subunit in mammalian cells where it functions to markedly potentiate transcriptional elongation^40, 41^. Interestingly, recent structural studies have suggested that the stimulation of Pol II elongation by RTF1 depends on its interaction with other components of the Pol II elongation complex particularly DSIF^41^. Notably, our CRISPR-ChIP screen identified both components of DSIF (SUPT4H1 and SUPT5), Iws1 (Supt6h), Tceb3 (Elongin A) and Ints3 (**Fig. 3a**), which are all known to function as potentiators of Pol II elongation^41–43^. Although, H3K79me2 has been loosely associated with transcriptional elongation, functional experimental evidence linking H3K79me2, and Pol II elongation has been scarce. Rather remarkably, our functional genomics approach identified the key subunits of the (i) PAF-complex^44^, (ii) DSIF complex^45^, (iii) Elongin complex^46^ and (iv) Integrator complex^47^, which have all been functionally validated to affect pause release and Pol II elongation rate^48, 49^. These results strongly support the notion that the maintenance of H3K79me2 by the DOT1L complex is dependent on efficient Pol II elongation.

As our CRISPR-ChIP approach was designed to enable us to uncouple the regulation of H3K79me2 from the requirement for malignant cell viability, we next wanted to assess which of the major regulators of H3K79me2 were also required for the survival of the MLL-AF9 leukaemia cells. As expected, many of the components required for Pol II elongation are common essential proteins and similarly, RNF20 and DOT1L which are established dependencies in MLL-AF9 leukaemia, also read out in our viability screen (**Fig. 3b and Supplementary Table 4**). Surprisingly, while MLLT10 is known to recruit DOT1L to chromatin via its PZP domain^50^ and clearly reads out as being required for H3K79me2 levels, it did not appear to be required for leukaemia cell survival (**Fig. 3b and Extended Data Fig. 3b**). To investigate these unexpected findings further we initially used two independent sgRNAs against each of the components of the DOT1L complex and assessed the effects on both global H3K79me2 levels and cell survival in MLL-AF9 leukaemia cells. These data confirmed that loss of DOT1L results in a global depletion of H3K79me2 and also dramatically reduces the survival of MLL-AF9 leukaemia cells^6, 51^ (**Fig. 3c-d**). Moreover, we also validated previous results showing that ENL is a dependency in MLL-AF9 leukaemia^52, 53^ but does not alter global H3K79methylation levels (**Fig. 3c-d**). Notably however, we confirmed that loss of MLLT10 has a profound effect on global H3K79me2 levels, but this is not accompanied by a survival disadvantage for the leukaemia cells (**Fig. 3c-d**). These findings are recapitulated in the Dep-Map database where all the MLL-leukaemia cell lines show a dependency on ENL for survival but only the cell lines, MOLM13 and MOLM14, both derived from the same patient, and SHI-1 (MLL-AF6), which does not aberrantly recruit DOT1L, show a dependency on MLLT10 (**Fig. 3e**). To explore the functional consequences of depletion of the four members of the DOT1L complex, we performed RNA-seq seq analysis following CRISPR/Cas9 deletion of each complex member (**Extended Data Fig. 3a-c**). When we analyse the top 100 genes downregulated by DOT1L loss, we also see a concomitant modest decrease in expression of these genes after depletion of ENL (**Fig. 3f**). Strikingly however, the genes most affected by DOT1L loss in MLL-AF9 leukaemia cells do not change their expression when either MLLT10 or MLLT6 are depleted (**Fig. 3f and Extended Data Fig. 3c-e**). Together, these surprising findings raise the prospect that although DOT1L is critical, global H3K79methylation levels maybe dispensable in MLL leukaemia and highlight our incomplete understanding of the regulation of H3K79methylation in this disease.

### Selective regulation of transcription by MLL-AF9

In order to address the discrepancy between H3K79me2 regulation and leukaemia cell survival by MLLT10 and DOT1L, we first sought to determine the influence of the MLLAF9 oncogene on both H3K79methylation and transcription of its direct target genes.

Current approaches for assessing the transcriptional influence of chromatin proteins largely rely on coupling methods to map chromatin occupancy (such as ChIP-seq) with changes in transcriptional output (using various RNA-Seq methods) following a perturbation of the target of interest ^1^. However, in the absence of small molecule inhibitors, the slow kinetics of genetic depletion experiments (e.g. shRNA/CRISPR) hampers the accurate assessment and separation of primary and secondary transcriptional changes. These limitations can be overcome by genetically engineered approaches that hijack the endogenous protein degradation machinery for rapid degradation of the target of interest by small molecules^54–56^. We adopted this system by generating a murine retroviral model of MLL-AF9 leukaemia engineered to contain a 3xFLAG tag for efficient mapping by ChIP-seq and a minimal auxin inducible degron (mAID) for rapid degradation using the plant hormone, auxin (IAA)^55^. To ensure that the MLL-AF9- F3-mAID construct faithfully recapitulated the disease biology induced by MLL-AF9, we transplanted our *in vitro* transformed haematopoietic stem and progenitor cells (HSPC) into C57/BL6 mice (**Fig. 4a**). We generated both MLL-AF9-F3-mAID and MLL-AF9-F3-mAID + mutant NRAS transformed cells and showed that both genotypes engrafted and gave rise to fulminant leukaemia with the expected disease latency (**Extended Data Fig. 4a**). To determine the genome-wide occupancy of MLL-AF9, we performed FLAG ChIP-seq in biological duplicates from MLL-AF9-F3-mAID along with ChIP-seq for Menin and H3K79me2 as well as a FLAG ChIP-seq from non-FLAG tagged MLL-AF9 transformed cells as a negative control (**Fig. 4b-c and Extended Data Fig. 4b-c**). The MLL-AF9-F3-mAID ChIP-seq data shows a high level of enrichment at previously reported MLL-AF9 target genes (e.g., Meis1 and Hoxa9) (**Fig. 4d and Extended Data Fig. 4b-c**). Notably, genes bound by MLL-AF9 were also co-occupied by Menin and H3K79me2, with all three factors displaying highly correlated enrichment (**Fig. 4c-d**).

**Figure 4:**
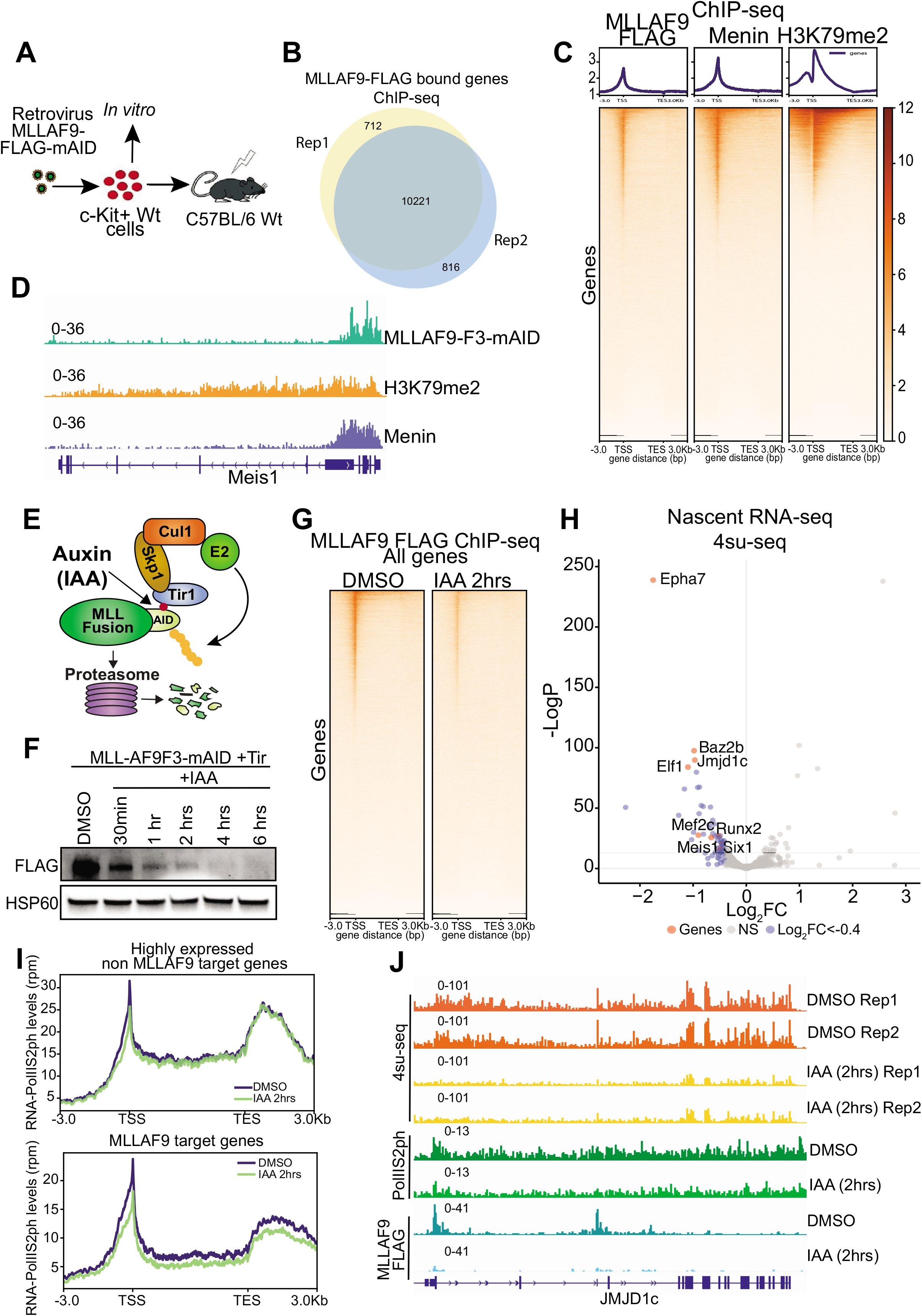
Selective regulation of transcription by MLL-AF9. **A)** Schematic of retroviral transformation of cKit+ HSPC with MLL-AF9 containing a c-terminal 3xFLAG and mAID tag. **B)** Venn diagram of MLL-AF9 FLAG ChIP-seq peaks from two replicates. **C)** Heatmap of MLL-AF9 FLAG, Menin, and H3K79me2 ChIP-seq in MLLAF9-F3-mAID cells. **D)** Genome browser snapshot of MLL-AF9, Menin and H3K79me2 at the *Meis1* locus. **E)** Schematic of the auxin-inducible degron system. **F)** Western blot analysis of MLL-AF9 degradation following an auxin (IAA) treatment timecourse using antibodies against FLAG and HSP60 (loading control). **G)** Heatmap of MLL-AF9 ChIP-seq in DMSO or IAA (2hrs) from MLLAF9-F3-mAID cells. **H)** Volcano plot of differentially expressed genes from nascent RNA-seq (4su-seq) in MLLAF9-F3-mAID cells treated with DMSO or IAA for 2hrs. Downregulated genes in blue (<LFC -0.4) and key direct target genes highlighted in red. **I)** Average profile plot of RNA-Pol IIS2ph ChIP-seq at highly expressed non-MLLAF9 and MLL-AF9 target genes in DMSO and IAA treated cells. **J)** Genome browser snapshot of the JMJD1c locus for MLL-AF9 and Pol IIser2 ChIP-seq and 4su-seq after 2hrs of DMSO or IAA treatment.

To determine the genes directly regulated by MLL-AF9, we treated MLL-AF9-F3-mAID cells expressing the plant Tir1 E3 ligase gene with indole-3-acetic acid (IAA) to facilitate IAA-dependent degradation of MLLAF9 (**Fig. 1e**). Time course analysis of MLL-AF9-F3-mAID + Tir1 cells treated with IAA showed near complete degradation within 2-4 hours of treatment (**Fig. 4f**). MLL-AF9 degradation resulted in a marked proliferation arrest and induction of differentiation, consistent with the role of MLL-AF9 in the maintenance of the leukaemic state^57^ (**Extended Data Fig. 4d-e**). To then identify the direct targets of MLL-AF9, we treated cells for 2hrs with IAA and performed FLAG ChIP-seq for MLLAF9-F3-mAID and nascent RNA-seq (4su-seq). FLAG ChIP-seq showed a dramatic global loss of MLL-AF9 from chromatin; however, nascent RNA-seq uncovered a core set of genes (Padj < 0.05) representing only ∼2% (260/10221) of the genes bound by MLL-AF9 that were immediately downregulated following MLL-AF9 degradation (**Fig. 4g-h and Extended Data Fig. 4f-h**). To complement the nascent RNA-seq data we also performed ChIP-seq for RNA polymerase II phosphorylated on serine 2 (RNA-PolIIS2ph). While we observed a global loss of MLL-AF9 from chromatin following 2hrs of degradation, global levels of RNA-PolIIS2ph were largely unchanged (**Extended Data Fig. 4i**). However, in agreement with the nascent RNA-seq data, we observed a selective loss of RNA-PolIIS2ph only at the MLL-AF9 bound genes that change their expression 2 hours after degradation of the oncoprotein (**Fig. 4i-j and Extended Data Fig. 4j**). Taken together, these data illustrate that the chromatin localisation of the oncoprotein is insufficient in isolation to assign a regulatory influence on transcription and emphasises the fact that although a large number of active genes are bound by MLL-AF9, only a small subset of these are fully reliant on the MLL-FP for their expression.

### MLL-AF9 selectively influences H3K79methylation at distinct target genes

Recently published data revealed that inhibition of either Menin or DOT1L with small molecule inhibitors effectively evicted MLL-AF9 from chromatin^56^ (**Fig. 5a**). However, the converse, which is the impact of global MLLAF9 eviction on H3K79methylation was not comprehensively assessed. Indeed, we confirmed the previous reports that both Menin (VTP50469) and DOT1L (SGC0946) inhibition caused the global eviction of MLL-AF9 from chromatin after 48 and 72hrs of treatment, respectively (**Fig. 5b, d and Extended Data Fig. 5a)**. Nonetheless, a major conundrum when assessing the chromatin localisation of transcriptional activators is that it is often hard to distinguish whether displacement of the regulator from chromatin is simply a consequence of a decrease in transcription. Despite, numerous studies assessing the transcriptional effects of MLL fusion proteins such as MLL- AF9, none have specifically addressed or answered this important issue. To resolve this issue, we decided to rapidly degrade the BET bromodomain proteins as targeting these proteins has previously been shown to be effective in MLL leukaemia cells^58, 59^ and rapid degradation leads to a profound and global loss of transcription^60, 61^. Remarkably, we found that even after 8 hours of treatment with the IBET-VHL degrader, MLL-AF9 remained bound to chromatin (**Fig. 5c, e and Extended Data Fig. 5b**). Collectively, these data emphasise two important previously unappreciated facts (i) MLL-FP are bound at chromatin in a manner that is insensitive to the transcriptional status of the locus and (ii) that both H3K79me2 and Menin are specifically important for the chromatin occupancy of MLL-FP.

**Figure 5:**
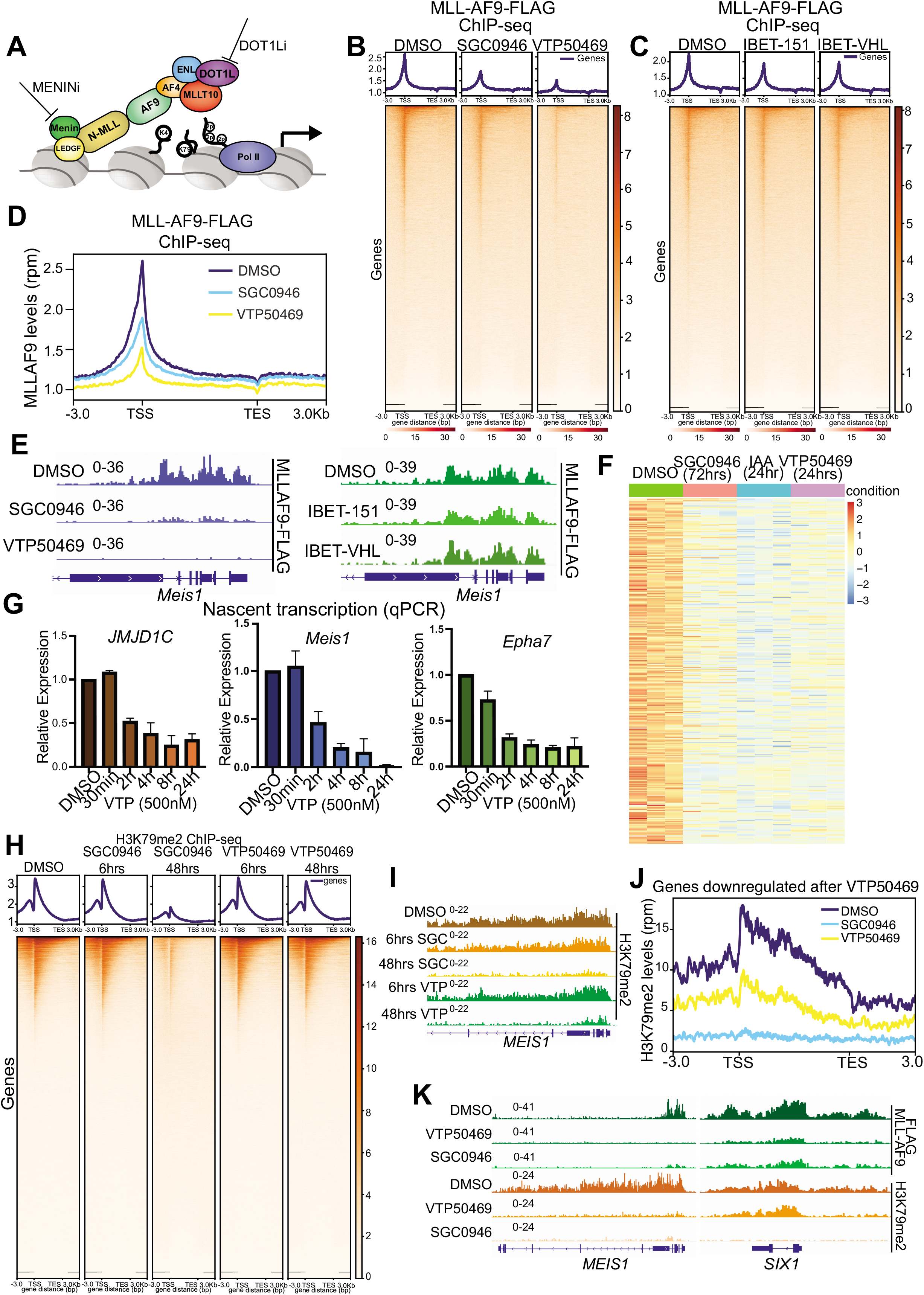
Selective regulation of H3K79me2 at target genes by MLL-AF9. **A)** Schematic of MLLAF9 oncogene with some of the associated cofactors along with their targeted therapies. **B)** Heatmap of MLLAF9 ChIP-seq after treatment with either DMSO, SGC0946 (5uM) for 72hrs, or VTP50469 (500nM) for 48hrs. **C)** Heatmap of MLLAF9 ChIP-seq after treatment with either DMSO, IBET-151 (500nM) for 8hrs, or IBET-VHL (500nM) for 8hrs. **D)** Average profile plot of MLLAF9 ChIP-seq across all genes after treatment with either DMSO, VTP50469 (500nM) and SGC0946 (5uM). **E)** Genome browser snapshots of the *Meis1* locus in MLLAF9-F3-mAID cells treated with DMSO, SGC0946, VTP50469 or IBET- 151 and IBET-VHL. **F)** Heatmap of downregulated genes from 3’RNA-seq in MLLAF9-F3- mAID cells treated with SGC0946 (72hrs), IAA (24hrs), or VTP50469 (24hrs). **G)** Nascent qPCR (primary transcript) analysis of VTP50469 timecourse treatment (500nM) vs DMSO in MLLAF9-F3-mAID cells using primers against *Jmjd1c*, *Meis1*, and *Epha7,* data normalised to housekeeping gene *(Gapdh),* error bars represent mean+/- SD from 3 biological replicates. **H)** Heatmap of H3K79me2 ChIP-seq analysis in MLLAF9-F3-mAID cells treated with either DMSO, SGC0946 (6hrs), SGC0946 (48hrs), VTP50469 (6hrs), and VTP50469 (48hrs). **I)** Genome browser snapshot of the Meis1 locus from the H3K79me2 ChIP-seq in (H). **J)** Average profile plot of H3K79me2 ChIP-seq across the body of genes downregulated after VTP50469 and bound by MLLAF9 in DMSO, SGC0946, or VTP50469 treatment. **K)** Genome browser snapshot of exemplar VTP50469 target genes with differential changes in H3K79me2 after VTP50469, *Meis1* (changed) and *Six1* (unchanged).

Whilst loss of H3K79me2 clearly resulted in a displacement of MLL-AF9, we and others have previously demonstrated that despite rapid enzymatic inhibition of DOT1L, H3K79me2 levels require several days to decrease^7, 62^. Therefore, it remained unclear what role H3K79me2 plays in the immediate early transcriptional regulation of MLL-AF9 target genes. Similarly, although Menin inhibition has recently been shown to unexpectedly result in a decrease in H3K79me2 levels, the molecular mechanism underpinning this observation is unknown^63^. To address these questions, we first sought to determine the transcriptional changes that are induced following treatment with SGC0946 and VTP50469 for 72hrs and 24hrs, respectively, on the core MLL- AF9 targets we identified. We chose these time points as this is when we and others observed efficient displacement of MLLAF9 from chromatin with these specific inhibitors of DOT1L and Menin^56^(**Fig. 5b, d**). We found that most of the genes repressed after MLL-AF9 degradation were also downregulated by both inhibitors, albeit at later time-points (**Fig. 5f and Extended Data Fig. 5c**).

Although the same genes were regulated by MLL-AF9 degradation, Menin and DOT1L inhibition, our data raised the prospect that the time frame in which these genes were altered with each of these interventions was substantially different. Given the slow turnover of H3K79me2 we considered the possibility that the levels of H3K79me2 alone are insufficient to maintain transcription of MLL-FP targets. To explore this further, we first wanted to determine the kinetics of Menin inhibition in relation to DOT1L inhibition at the same genes. We performed primary transcript-qPCR (nascent qPCR) after a time course with VTP50469 treatment for up to 24hrs and found that Menin inhibition resulted in rapid downregulation of key targets (*Meis1, Jmjd1c, Epha7*) as early as 2 hrs, which closely paralleled our findings with MLL-AF9 degradation (**Fig. 5g**). To understand how this is reflected at chromatin, we performed ChIP-seq for H3K79me2 and MLLAF9-FLAG after early and late treatment with VTP50469 and SGC0946. Consistent with our previous studies using SGC0946, H3K79me2 was only reduced after 48hrs of treatment with no global changes detected after 6hrs (**Fig. 5h- i**). Importantly, ChIP-seq analysis shows eviction of MLL-AF9 after 6 and 48hrs of VTP50469 treatment with no global changes in H3K79me2 (**Fig. 5h-i and Extended Data Fig. 5d**). These results suggest that transcription of key MLL-FP target genes can be markedly suppressed without any change in H3K79me2.

Interestingly, although global H3K79me2 levels are not changed with Menin inhibition at 6 hours or 48hrs (**Fig. 5h**), we observed a potent suppression of H3K79me2 after 48 hrs of VTP50469 treatment at key MLL-target genes (**Fig. 5i**), raising the prospect that the Menin-MLL-Fusion complex recruits DOT1L to chromatin to deposit H3K79me2 at a small subset of genes. Notably, when we looked at the genes where VTP50469 treatment results in transcriptional repression, we find that MLL-AF9 displacement is accompanied by a profound loss of H3K79me2 (**Fig. 5j and Extended Data Fig. 5e-f**). However, even though VTP50469 results in a global displacement of MLL-AF9 (**Fig. 5b, d**) not all MLL-AF9 bound genes show a loss of H3K79me2 after displacement of the oncoprotein with Menin inhibition (**Fig. 5k and Extended Data Fig. 5e-f**). Taken together, these results suggested a multifactorial regulation of H3K79methylation in MLL-FP leukemia which cannot be solely explained by the simple model of aberrant recruitment of DOT1L by MLL-AF9.

### MLLT10 regulates H3K79methylation at regions not occupied by MLL-FP

The selective effects of Menin inhibition and MLLAF9 eviction on H3K79methylation alongside the clear requirement for Menin and MLL-AF9 in the maintenance of leukaemia cells survival, sharply contrasted with the negligible effects on leukaemia cell survival after loss of MLLT10, despite there being a profound global loss of H3K79me2 levels.

To explore the mechanistic basis for this observation, we performed spike-in normalised ChIP-seq for H3K79me2 after depletion of DOT1L or MLLT10 with two independent sgRNAs. These data provided orthogonal confirmation that both MLLT10 and DOT1L regulate H3K79me2 levels globally (**Extended Data Fig. 6a**). Notably, when we look specifically at our previously annotated MLL-AF9 target genes (**Fig. 4h**), we find that H3K79me2 levels are higher at these genes compared with Non-MLL-AF9 target genes (**Fig. 6a-b**). More surprising is that the fact that whilst H3K79me2 levels are markedly reduced after DOT1L loss, there are no changes in H3K79me2 levels at the major MLL-AF9 target genes including MEIS1 and HOXA9 following MLLT10 loss (**Fig. 6a-c and Extended Data Fig. 6b-c**). Together these findings raised the intriguing prospect that in MLL-AF9 leukaemia, DOT1L is functionally partitioned into two separate complexes; one that contains the malignant MLL-AF9 oncogene and one that is the native endogenous DOT1L complex containing MLLT10.

**Figure 6:**
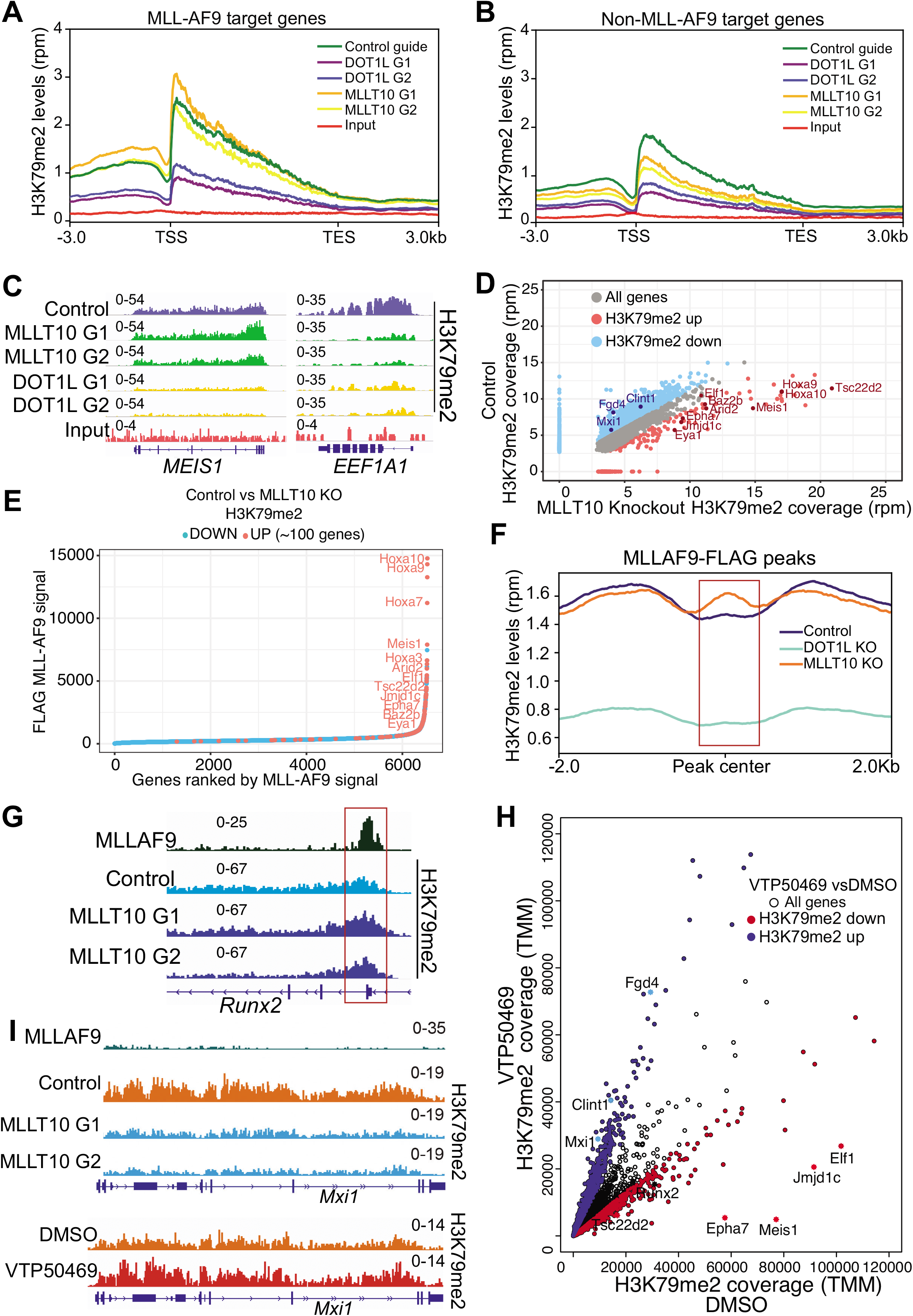
DOT1L functions in distinct native and oncogenic complexes in MLL-FP leukaemia. **A)** Average profile plot of H3K79me2 levels at MLL-AF9 target genes in MLL-AF9^FLAG-mAID^ Cas9 cells transduced with either a non-silencing guide (Control) or 2 independent guides targeting DOT1L (G1 and G2) or MLLT10 (G1 and G2). **B)** Average profile plot of H3K79me2 levels at the top 500 expressed genes <u>not</u> regulated by MLL-AF9 in MLL-AF9^FLAG-mAID^ Cas9 cells transduced with either a non-silencing guide (Control) or 2 independent guides targeting DOT1L (G1 and G2) or MLLT10 (G1 and G2). **C)** Genome browser snapshots of canonical MLL-AF9 target gene *Meis1* and the *Ef1a* locus used in the CRISPR-ChIP plasmid. Tracks show levels of H3K79me2 and input in MLL-AF9 Cas9 cells transduced with either a non- silencing guide (Control) or two independent guides targeting DOT1L or MLLT10. **D)** Correlation plot of H3K79me2 ChIP-seq peak size in non-silencing control vs MLLT10 knockout (average of two independent guides) showing genes with increased H3K79me2 (red) or decreased (blue). **E)** Hockey stick plot of genes ranked by ^FLAG^MLL-AF9 occupancy; red dots represent genes that display increased H3K79me2 following MLLT10 KO vs non- silencing guides (Control) while blue dots represent genes with decreased H3K79me2 following MLLT10 KO. **F)** Profile plot of H3K79me2 distribution across MLLAF9 peaks +/- 2kb in MLLAF9^FLAG-mAID^ Cas9 cells transduced with control guides or guides targeting DOT1L or MLLT10. Red box over the MLL-AF9 peak. **G)** Genome browser snapshot of exemplar direct and strongly bound MLL-AF9 target gene that shows increased H3K79me2 following MLLT10 KO in MLL-AF9 cells, *Runx2*. Red box over the MLL-AF9 peak. **H)** Correlation plot of H3K79me2 normalised count coverage in vehicle treated (DMSO) or Menin inhibitor (VTP50469) murine MLL-AF9 cells. Genes with a decrease in H3K79me2 following VTP50469 are highlighted in red and genes with increased H3K79me2 are in blue. **I)** Genome browser snapshot of an exemplar Menin inhibitor non-responsive gene, *Mxi1*, with no MLL- AF9 occupancy, showing increased H3K79me2 after VTP50469 treatment and decreased H3K79me2 in MLLT10 KO cells compared to control cells.

The possibility of two distinct DOT1L complexes within MLL-AF9 leukaemia cells was further supported by the fact that disruption of MLLT10 in the leukaemia cells appeared to increase H3K79me2 levels at known MLL-AF9 target genes (**Fig. 6c**). To explore this further we looked globally to identify genes where H3K79me2 levels were unchanged or increased following MLLT10 depletion. These data clearly identified ∼200 genes where H3K79me2 levels were elevated rather than depleted following MLLT10 loss (**Fig. 6d**). Remarkably, when we assess the binding of MLL-AF9 across the genome we find that the genes with the greatest chromatin occupancy of MLL-AF9 are the genes where H3K79me2 levels are increased following MLLT10 loss (**Fig. 6e**). In contrast, H3K79me2 levels decrease after MLLT10 depletion at the genes with the lowest occupancy of MLL-AF9 (**Fig. 6b, e**). Together these data suggest that loss of MLLT10 results in the disassembly of the native DOT1L complex containing MLLT10, which in turn results in a relative excess of DOT1L that is able to associate with MLL-AF9 and further accentuate H3K79me2 levels at the major target genes of the oncogenic complex. In further support of this contention, is the striking observation that H3K79me2 was selectively increased directly where MLLAF9 is bound but was reduced in the gene body of MLLAF9 target genes following loss of MLLT10 (**Fig. 6f-g and Extended Data Fig. 6d**).

To further strengthen these findings, we sought to demonstrate the existence of biochemically distinct complexes using size exclusion chromatography. Whilst our use of an epitope tag to MLL-AF9 enabled our detailed studies of chromatin binding in mouse cells, there are unfortunately no reliable antibody reagents to detect mouse MLLT10, DOT1L, ENL and MLLT6. However, as many of the paradigms established for MLL leukaemias in mouse cells are conserved in human cells and moreover shared by several of the common MLL-fusions we chose to biochemically characterise cellular extracts from the most commonly used human MLL-leukaemia cell line MV4;11 containing an MLL-AF4 fusion. When analysed by western blot these data showed that although MLLT10 and DOT1L co-elute (fraction 3), DOT1L but not MLLT10 co-elutes with MLL1 (**Extended Data Fig. 6f**). Taken together with the ChIP-seq analyses these findings are consistent with the existence of two distinct DOT1L complexes in MLL-FP leukaemia: (i) an oncogenic DOT1L complex containing the MLL-FP and (ii) a native DOT1L complex containing MLLT10.

Our earlier data suggested that whilst Menin inhibition has no influence on the regulation of H3K79me2 by the native endogenous DOT1L complex which includes MLLT10, it does dramatically affect DOT1L activity as it displaces the oncogenic DOT1L complex associated with MLL-AF9 from chromatin (**Fig. 5b, j**). Consistent with the fact that there is a reciprocal re-distribution of DOT1L between the oncogenic and endogenous complexes, we find that displacement of chromatin bound MLL-AF9 after Menin inhibition results in an increase in H3K79me2 at genes dependent on the endogenous MLLT10/DOT1L complex (**Fig. 6h-i**). These results further support the fact that DOT1L exists in both a native MLLT10-dependent complex and an MLL-AF9 oncogenic complex, which together cooperate to regulate H3K79methylation at critical MLL-FP target genes. These data also provide a molecular explanation for the differential regulation of H3K79me2 by Menin inhibition at MLL-AF9 target genes (**Fig. 5i-k and Extended Data Fig. 5e-f**).

### Native and oncogenic DOT1L complexes cooperate to regulate MLL-FP target genes

To demonstrate the potential clinical implications of the partitioning of DOT1L into two functional complexes, we assessed how abolition of the native complex (through MLLT10 loss) impacts the transcriptional response of MLLAF9 cells to VTP50469. We treated control or MLLT10 depleted MLLAF9 cells with a low dose of Menin inhibitor (VTP50469 20nM) or DMSO for 48hrs and performed RNA-seq analysis. When we looked at target genes directly bound and regulated by MLLAF9 (including *Meis1*, *Six1*, and *Epha7*), we found that these genes were substantially more downregulated by Menin-inhibition in the MLLT10 depleted cells compared to control cells (**Fig. 7a**). Indeed, the majority of the Menin/MLLAF9 target genes showed a more pronounced downregulation in MLLT10 depleted cells after VTP50469 treatment (**Fig. 7b and Extended Data Fig. 7a**). We next explored whether these transcriptional responses would translate to a functional outcome by performing proliferation assays in the MLLT10 depleted vs control cells treated with VTP50469. These data show that whilst cells deficient in MLLT10 have no growth disadvantage in the absence of drug (**Extended Data Fig. 7b**), the resulting absence of a native DOT1L complex, sensitises the cells to Menin inhibitors (**Fig. 7c and Extended Data Fig. 7b**). To provide a mechanistic explanation for this, we looked at H3K79methylation by ChIP-seq and observed that in the absence of the native DOT1L complex (deletion of MLLT10), H3K79me2 levels were markedly more sensitive to treatment with VTP50469 compared to control cells at MLL-FP target genes (**Fig. 7d and Extended Data Fig. 7c**). These results indicate that specific disruption of the native DOT1L complex causes H3K79me2 at MLL-FP targets to be largely maintained by the oncogenic DOT1L complex, which substantially enhances the transcriptional response and sensitivity to Menin inhibition.

**Figure 7:**
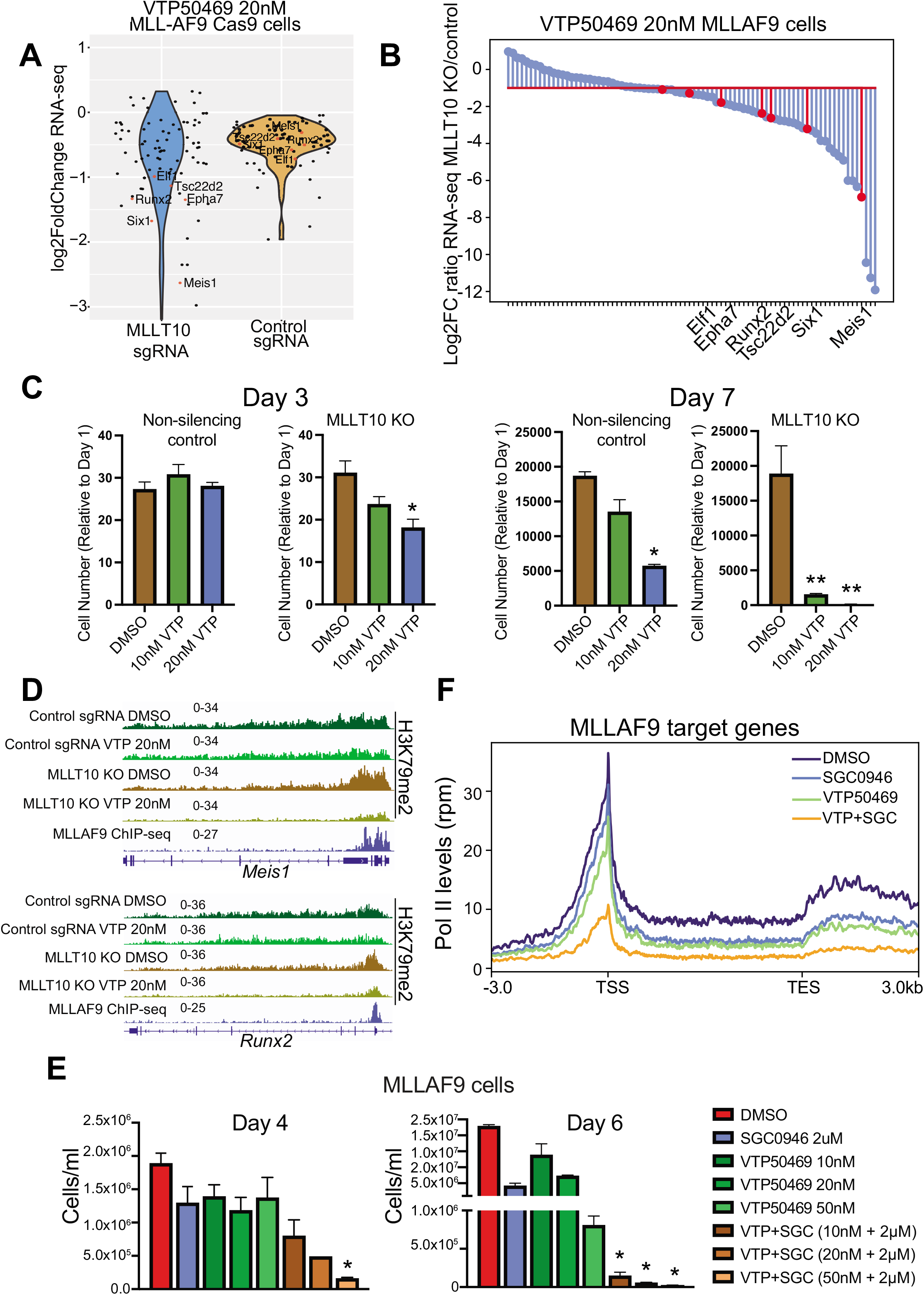
Disruption of the native DOT1L complex sensitises MLLAF9 leukaemia cells to Menin inhibition. **A)** Violin plot of RNA-seq in MLLAF9^FLAG-mAID^ Cas9 cells expressing control guides or MLLT10 guides (stable knockout of MLLT10) and treated with either DMSO or VTP50469 (20nM). Menin inhibitor target genes that are also bound by MLLAF9 are shown. **B)** Lollipop plot of RNA-seq logFC of the ratio of Control vs MLLT10 KO after treatment with VTP50469 (20nM) in MLLAF9^FLAG-mAID^ cells. **C)** Proliferation assay in control and MLLT10 KO MLLAF9^FLAG-mAID^ Cas9 cells treated with either DMSO, 20nM or 40nM VTP50469. Cell counts shown at day 3 and day 7 and normalised to day 1 counts. n=3 biological replicates. * p <0.05 ** p<0.01. **D)** Genome browser snapshot of exemplar MLLAF9 targets *Meis1* and *Runx2* from H3K79me2 ChIP-seq in MLLAF9^FLAG-mAID^ cells with control guides or MLLT10 KO and treated with vehicle (DMSO) or VTP50469 (20nM) for 48hrs. MLLAF9 tracks shown for reference. **E)** Proliferation assay in MLLAF9^FLAG-mAID^ Cas9 cells treated with either DMSO, SGC0946 (2uM), VTP50469 (10nM, 20nM or 50nM) and combinations. Cell counts shown at day 4 and day 6 and normalised to day 1 counts. n=3 biological replicates. * p <0.05. **E)** Profile plot of RNA-Pol II ChIP-seq enrichment at MLLAF9 target genes after treatment with either DMSO, SGC0946, VTP50469 or Combo for 48hrs.

Since there are currently no small molecules available against MLLT10, we instead used DOT1L inhibitors to determine the potential synergy of targeting the Menin-dependent oncogenic DOT1L complex with Menin inhibition alongside inhibitors of the native DOT1L complex. We demonstrate that combination of DOT1L and Menin inhibitors showed strong synergistic efficacy (**Fig. 7e and Extended Data Fig. 7d-e**). Consistent with the greater functional inhibition of MLL-AF9 cells we also noted that the levels of chromatin bound RNA- Pol II at MLL-AF9 target genes were more dramatically reduced in the combination treated cells (**Fig. 7f)**. Taken together, these data highlight that both the oncogenic and native DOT1L complexes cooperate in the maintenance of MLL-FP target gene expression and provide the molecular rationale for potential future combination therapy strategies using DOT1L and Menin inhibitors in MLL-FP leukaemia (**Fig. 8**).

**Figure 8:**
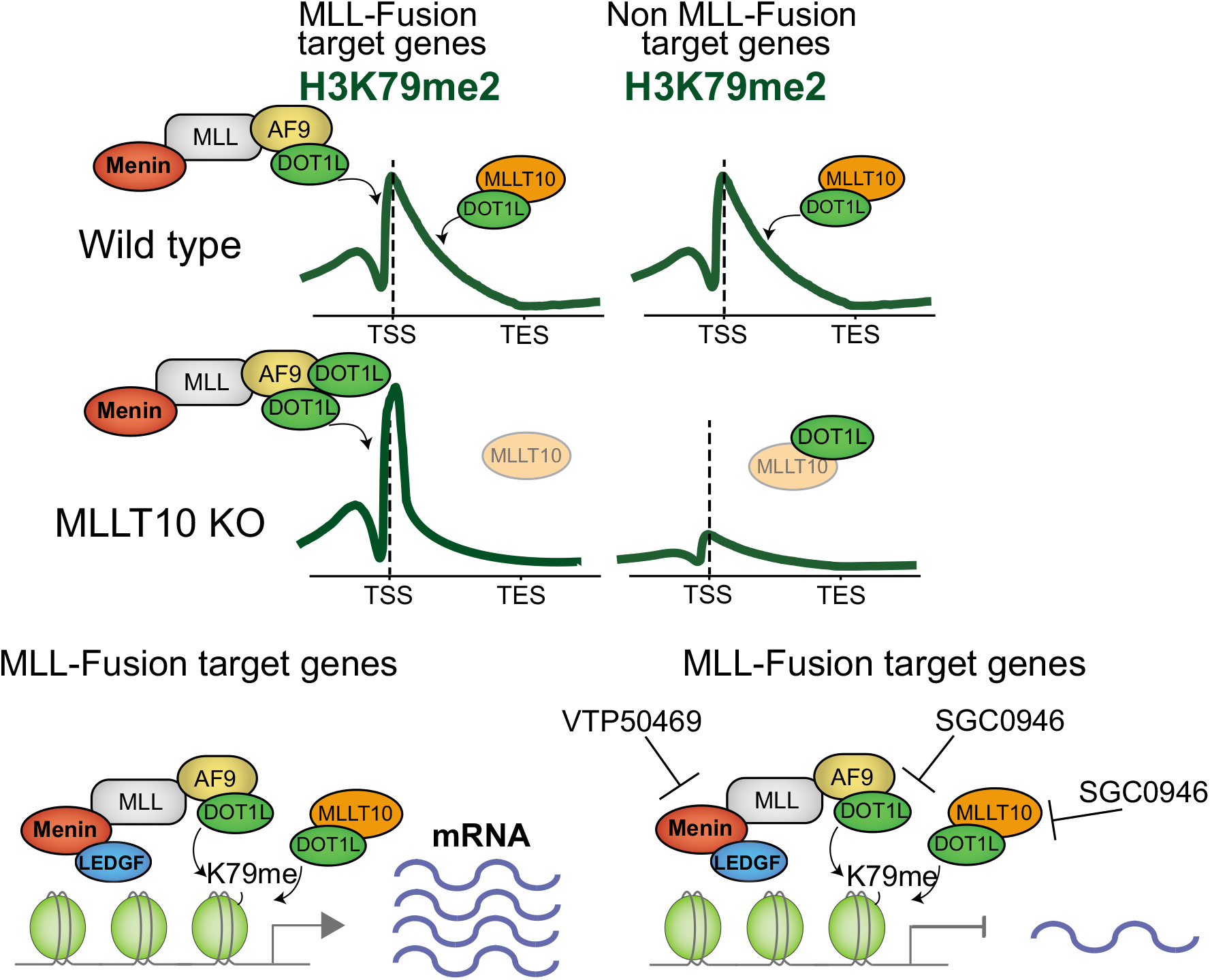
Schematic to illustrate the distinct function of the native and oncogenic DOT1L complex.

## Discussion

Despite being one of the most intensely studied subset of oncoproteins, the molecular mechanisms by which MLL-FP and its cooperating oncogenic cofactors regulate their target genes remain incompletely understood. Our study sought to address this key issue by focussing on the molecular regulation of a histone modification, H3K79me2, which has been a central focus of the oncogenic process mediated by MLL fusion proteins. Identifying the major regulatory proteins required for histone modifications originally involved a candidate approach which was markedly facilitated by yeast genetics coupled to labour-intensive biochemistry^64^. However, these biochemical strategies alone are unable to assign a functional role to complex members or identify independent functional regulators. As the cornerstone method for understanding chromatin biology has been ChIP^65^, we sought to couple ChIP to high-throughput CRISPR screens to enable a strategy for unbiased discovery of chromatin templated processes. Whilst our CRISPR-ChIP approach can be adapted to study any defined chromatin event, as a proof-of-concept, here we have used it to identify all of the major non-redundant regulators of two highly conserved histone modifications, H3K4me3 and H3K79me2. Using this novel method, we showed that each member of the WRAD complex is essential for efficient deposition of H3K4me3 by the COMPASS and COMPASS-like complexes. CRISPR-ChIP further allowed us to reveal the specific COMPASS complex (SET1A/B) that is likely functioning at the EF1a locus and that Pol II is dispensable for the maintenance of H3K4me3 genome-wide in human cells. We subsequently used CRISPR-ChIP to identify MLLT10 and DOT1L as key regulators of H3K79me2 genome-wide. Notably, our unique approach also uncovered an unexpected discrepancy between global H3K79me2 levels and leukaemia cell survival. The prevailing paradigm had suggested that H3K79me2 levels and leukemia cell viability were inexorably linked; however, our unbiased screens had illustrated that global levels of H3K79me2 can be profoundly diminished with loss of MLLT10 with a negligible influence on leukaemia cell survival. We reconcile these observations by illustrating that DOT1L exists in two functionally distinct complexes in MLL-FP leukaemia. The native complex includes MLLT10 which facilitates H3K79 methylation at the majority of transcriptionally active genes but has little influence on the maintenance of the malignant state. In contrast, DOT1L also exists in a neomorphic oncogenic complex with the MLL-FP, which is critical for the maintenance of the malignant transcription program and viability of the leukaemia cells.

Beyond establishing the molecular regulation of H3K79me2 levels in MLL-leukaemia our data also sought to address another contentious and unresolved issue in the field - is H3K79me2 necessary for transcription? Using a combination of approaches to accurately map chromatin occupancy of the MLL-AF9 oncogene and the subsequent rapid and selective degradation of the oncogene coupled to nascent transcriptomics we illustrate several key principles. (i) Although MLL-AF9 is bound to several thousand genes, only ∼2% of these genes have a discernible immediate and early decrease in transcriptional output following loss of the oncogene. (ii) The process of transcription per se does not influence the chromatin occupancy of MLL-FP, instead, the chromatin occupancy of this oncogenic fusion is selectively dependent on its interaction with Menin. In this regard, a key insight that our data provides relates to the different kinetics of transcriptional regulation between catalytic inhibition of DOT1L and disruption of Menin-MLL1 interaction. While MLLAF9 degradation and Menin inhibition result in rapid transcriptional repression, DOT1L inhibition takes days to alter transcription. The implications of these findings are important as they illustrate that although H3K79me2 is associated with transcriptional activity, it is not sufficient to maintain transcription and H3K79me2 is not the major regulator of MLL-FP target gene expression. Instead, our data suggests that H3K79me2 is likely required to maintain a chromatin state that is permissive for MLL-AF9 recruitment and binding.

Finally, our data highlighting the existence of native and oncogenic DOT1L complex provides the molecular explanation for the unexplained observation that Menin inhibition unexpectedly results in a delayed decrease in H3K79me2 at MLL-FP target genes. It also explains why inhibition of Menin in MLL-FP leukaemia results in an increase in gene expression and H3K79me2 levels at genes not bound by Menin or MLL-FP. Understanding how perturbation of one chromatin bound complex results in the dynamic flux of key members between sub- stoichiometric complexes leading to differential gene expression is an important emerging theme in chromatin regulation. Moreover, our findings of functionally distinct DOT1L complexes specifies the molecular rationale for observations of synergy between DOT1L and Menin inhibitors^66^. Taken together, they highlight why targeting both the endogenous and oncogenic DOT1L complexes with combination therapies may increase the therapeutic efficacy and clinical outcomes with these novel therapies against MLL-FP driven leukaemia.

## Acknowledgements

We thank the Flow Cytometry facility and Molecular Genomics Core at the Peter MacCallum Cancer Centre and the ARAFlowcore Flow facility at the Australian Centre for Blood Diseases, Monash University. We thank the following funders for fellowship, scholarship and grant support: VCA Mid-Career Research Fellowship (O.G); Cancer Council Victoria Sir Edward Dunlop Research Fellowship, NHMRC Investigator Grant #1196749 and Howard Hughes Medical Institute International Research Scholarship #55008729 (M.A.D); and NHMRC Project Grants #1146192 (O.G), #1085015 / #1106444 (M.A.D) and #1128984 (M.A.D).

## Author contributions

O.G and M.A.D conceived, designed and supervised the research and wrote the manuscript. O.G, C.C.B, D.N, D.F, K.K, M.B and C.D conducted experiments and/or analysed data. O.G, C.C.B and K.K performed the CRISPR-ChIP screens. E.Y.N.L and L.T led the analysis of the genomic data and CRISPR-ChIP screens, with contribution from Y-C.C.

## Data and code availability

Sequencing data has been deposited into the sequence read archive, hosted by the National Center for Biotechnology Information. The accession number for the sequencing data reported in this paper is NCBI sequence read archive: GSE192562.

## Materials and Methods

### Cell lines

K562, HEK293T, MOLM-13, MV4;11 and drosophila S2 cells were obtained from ATCC. MLL-AF9 leukaemic blasts were generated by magnetic bead selection (Miltenyi Biotec) of c- KIT^+^ cells from whole mouse bone marrow and subsequent retroviral transduction with an MSCV-FLAGx3-MLL-AF9 construct. K562, MV4;11 and MOLM-13 cells were cultured in RPMI-1640 supplemented with 2mM Glutamax, 100 IU/mL Penicillin, 100 μg/mL Streptomycin and 10% HI-FCS. HEK293T cells were cultured in DMEM supplemented with 2mM Glutamax, 100 IU/mL Penicillin, 100 μg/mL Streptomycin and 10% HI-FCS. MLL-AF9 cells were cultured in RPMI-1640 supplemented with 2 mM Glutamax, 100 IU/mL Penicillin, 100 μg/mL Streptomycin, 10% HI-FCS, IL-3 10 ng/mL, IL-6 10 ng/mL and SCF 50 ng/mL (Peprotech) for the first two weeks and then IL-3 10ng/mL for ongoing culturing. Drosophila S2 cells were cultured in Schneider’s Drosophila media (Life Technologies) supplemented with 100 IU/mL Penicillin, 100 μg/mL Streptomycin and 10% HI-FCS. All cell lines were cultured in 5% CO2 at 37°C with the exception of Drosophila S2, which was cultured at room temperature. All cell lines were subjected to regular mycoplasma testing and underwent short tandem repeat (STR) profiling.

### Chemicals

SGC0946 was obtained from structural genome consortium, VTP50469 and A-485 were purchased from Selleck chemicals. For each inhibitor doses used, and durations of treatment are indicated in figure legends. For ChIP-qPCR, A-485 was used at a dose of 3uM for 24hrs. For ChIP-seq experiments, VTP50469 was used at a dose of 500nM and cells typically harvested at 48hrs unless otherwise indicated. SGC0946 was used at a dose of 5uM for 72hrs. DMSO was used as vehicle control for all drug treatment expeirments.

### CRISPR sgRNA library

The targeted sgRNA CRISPR library is a custom built ∼7239 human and ∼7395 mouse sgRNAs Targeting ∼1144 chormatin regulators (6sgRNAs per gene), as well as non-targeting and safe targeting control sgRNAs (guide sequences are provided in Table S1). Library sgRNAs were expressed in the CRISPR-ChIP plasmid, which is a modified pKLV-U6gRNA(BbsI)- EF1apuro2ABFP lentiviral sgRNA expression vector, which encodes puromycin and BFP selection markers obtained from Addgene #50946, a gift from Kosuke Yusa ^67^ as described. Murine *Meis1* promoter comprised of ∼1.4kb upstream of TSS Murine *Myc* enhancer was obtained from H3K27ac ChIP-seq and ATAC-seq data ^68^ and cloned upstream of a minimal CMV promoter. 500bp of human PGK promoter and 1kb of human EF1a promoter including 5’UTR intron. The cis-elements were cloned into the PKLV guide plasmid ∼40bp downstream of the guide sequence.

### CRISPR-ChIP screens

K562 and MLLAF9 cells were transduced with a lentiviral vector encoding Cas9-IRES mCherry. mCherry positive cells were sorted as single cells to obtain a clonal population of high cas9 expressing cells.

Cas9 cells (>100 x 10^6^ per cell line) were infected with the pooled lentiviral sgRNA library at a multiplicity of infection of 0.3. The percentage of cells infected was determined by flow cytometry-based evaluation of BFP^+^ (sgRNA-expressing) cells 48 hours following transduction of the targeted epigenetic library. Infected cells were selected with 2 μg/mL puromycin commencing 48 hours after transduction. Viable cells were collected and sorted at day 6 (Pol II, H3K4me3) and day 9 (H3K79me2) for ChIP as well as library control (input) samples. For the ChIP samples, ∼120x10^6^ cells were cross linked with 1% formaldehyde (Sigma-aldrich). For the library control samples ∼5-10x10^6^ cells were harvested for genomic DNA. Lysed cells for ChIP were sonicated to obtain average fragment size of 1-2kb to ensure gRNA remains linked to promoter sequence. ChIP was performed as previously described, with some modifications, ∼8-10x10^6^ cells’ worth of chromatin was incubated with 2-4ug of antibody in separate tubes, between 12-15 ChIP tubes per CRISPR-ChIP experiment was used. 50ul of Protein A or G beads was added to each tube after O/N incubation followed by washing and elution. Elution was performed twice and de-crosslinked O/N. ∼12 ChIP tubes were purified using Qiagen PCR purification columns. 3 ChIP tubes were purified across one PCR cleanup column and column elution was performed at least 4 times in 50ul volume with heated nuclease-free water to maximise the yield of ChIP DNA.

One-step PCR was performed to amplify the guide sequence from ChIP DNA or genomic DNA, containing adaptors for Illumina sequencing. Samples were sequenced with single-end 75 bp reads on an Illumina NextSeq500. The sequence reads were trimmed to remove the constant portion of the sgRNA sequences with fastx clipper (http://hannonlab.cshl.edu/fastx_toolkit/) or cutadapt v1.14^69^, then mapped to the reference sgRNA library with bowtie2^70^. After filtering to remove multi-aligning reads, the read counts were computed for each sgRNA. MAGECK was used to rank the genes for which targeting sgRNAs were significantly enriched in the ChIP samples compared to the control samples taken from the same pool of growing mutagenized cells. MAGECK analysis of CRISPR-ChIP screens is in the Supplementary Tables 2-3, and CRISPR-dropout analysis is in Supplementary Table 4.

### CRISPR/Cas9-mediated gene disruption

Single guide RNA (sgRNA) oligonucleotides (Integrated DNA Technologies) were cloned into lentiviral expression vectors pKLV-U6gRNA(BbsI)-PGKpuro2ABFP (Addgene #50946, a gift from Kosuke Yusa)^67^ as described. Oligonucleotide sequences are listed in Table 2, with sources as indicated^71, 72^. For CRISPR/Cas9-mediated gene disruption, cells were first transduced with the Cas9 expression vector pHRSIN-PSFFV-Cas9-PPGK-Blasticidin^73^ or FUCas9Cherry (a gift from Marco Herold, Addgene #70182)^74^, and selected with blasticidin or sorted for mCherry expression respectively. To generate polyclonal populations with targeted gene disruption, cells were subsequently transduced with pKLV-gRNA- PGKpuro2ABFP encoding either gene-specific sgRNAs or with a control sgRNA targeting a ‘safe’ genomic location with no annotated function^71^. Cells infected with pKLV-gRNA- PGKpuro2ABFP were selected with 2 μg/mL puromycin for 72 hours, commencing 48 hours after transduction. Efficient functional CRISPR/Cas9-mediated gene disruption of target genes was confirmed by immunoblot or sanger sequencing using TIDE analysis.

**Table 1:**
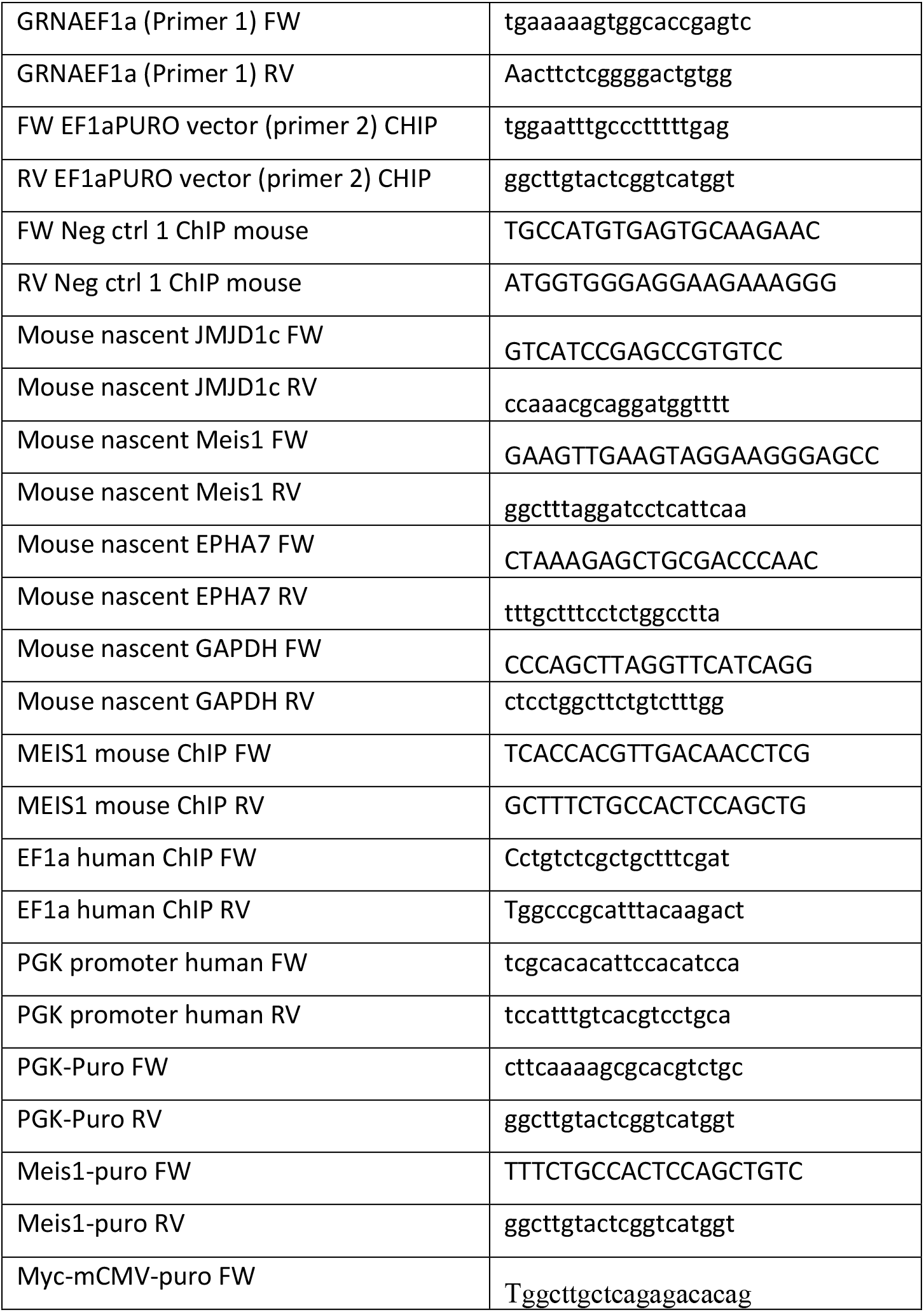
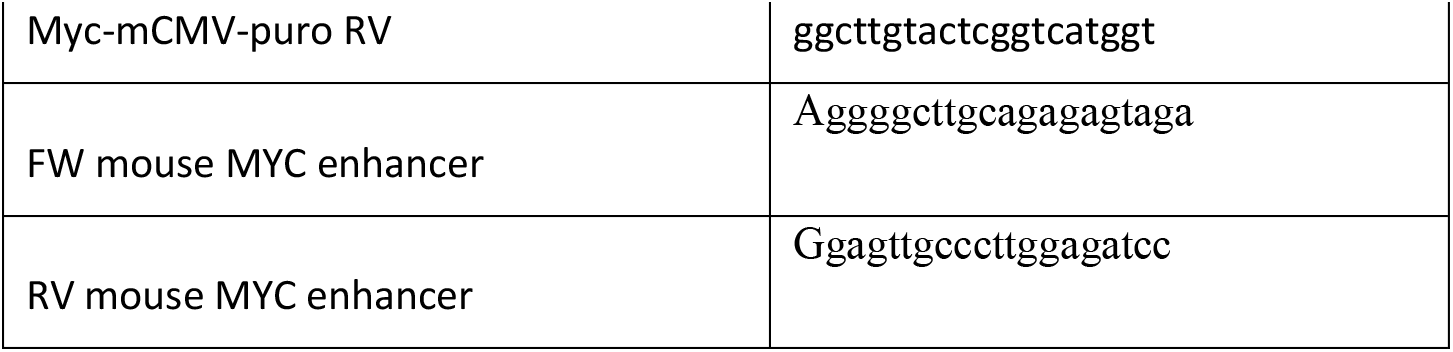
ChIP-qPCR and Nascent-qRT-PCR primers.

**Table 2:**
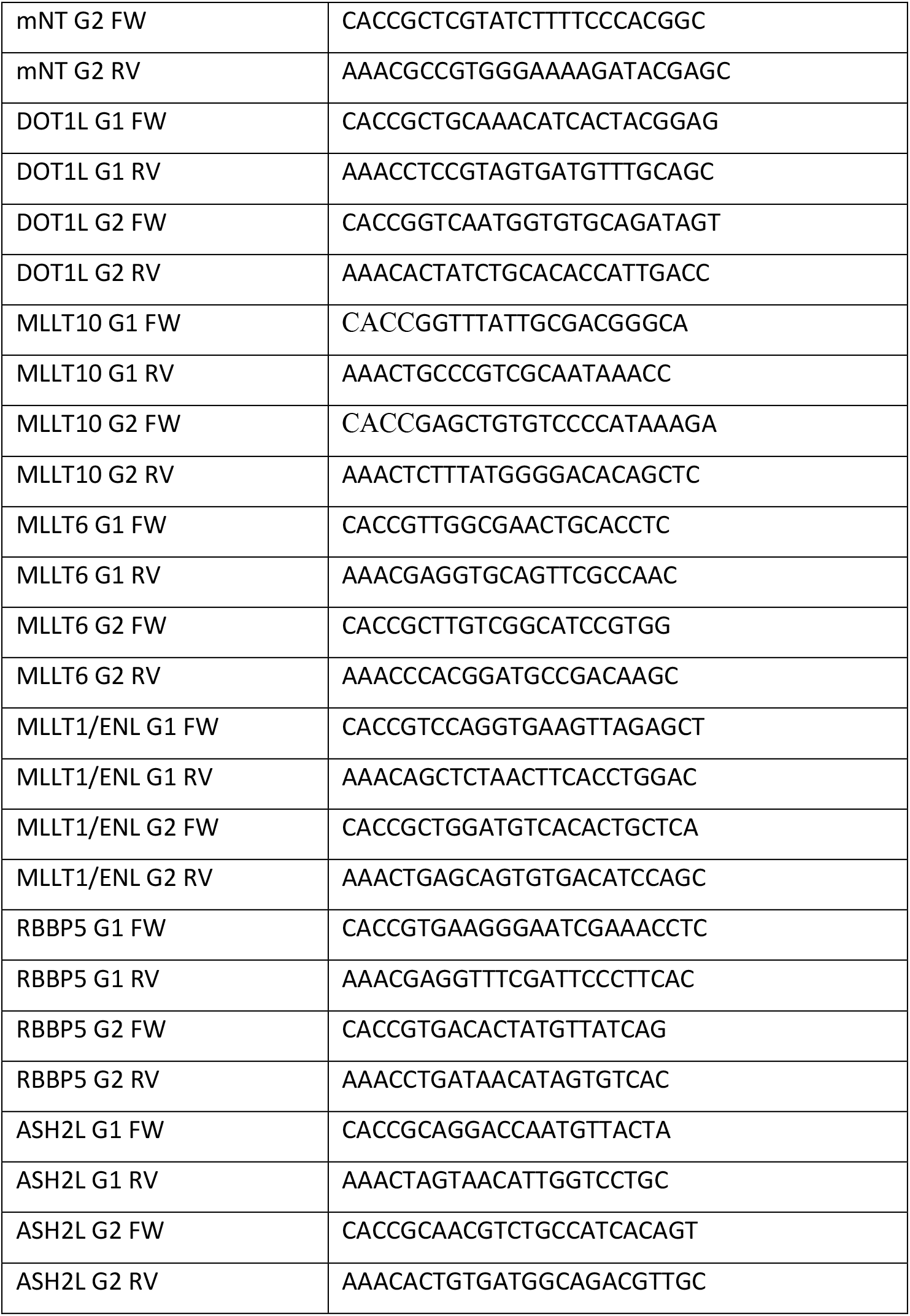
sgRNA oligos.

### Virus production and transduction

Lentivirus was produced by triple transfection of HEK-293ET cells with a lentiviral transfer vector and the packaging plasmids psPAX2 and pMD.G at a 0.5:0.35:0.15 ratio. Retrovirus was produced by triple transfection of HEK-293ET cells with a retroviral transfer vector, structural pMD1-gag-pol plasmid and pMD.G envelope plasmid at a 0.75:0.22:0.03 ratio. All transfections were performed using polyethylenimine (PEI). Viral supernatants were collected 48 hours following transfection, filtered through a 0.45 μm filter and added to target cells.

### Flow cytometry

Cells were washed in PBS/2% FCS, samples were resuspended in PBS with 2% FCS and filtered. Data were acquired on a BD LSRFortessa or BD FACSymphony and analysed in FlowJo or sorted using a BD FACSFusion sorter.

### Immunoblotting

Cells were lysed in 1% SDS in 100 mM Tris-HCl pH 8.0 with Roche cOmplete EDTA-free protease inhibitor at room temperature. DNA was fragmented by adding 1:100 Benzonase (Sigma). Lysates were heated to 70°C in SDS sample buffer with 50 mM DTT for 10 minutes, separated by SDS-PAGE and transferred to PVDF membrane (Millipore). Membranes were blocked in 5% milk in TBS-T and incubated with indicated antibodies O/N at 4 degrees. Blots were imaged with ECL prime using a gel doc instrument.

### Size exclusion chromatography

MV411 cell extract from 50 x 10^6^ cells in 500 µL was clarified by centrifugation at 21,000 g at 4°C for 10 min. 500 µL sample was then injected into a 500 µL loop and was loaded onto a Superose 6 Increase 10/300 GL column (Cytiva) equilibrated with Buffer A (50 mM HEPES pH 7.5, 250 mM NaCl, 10% glycerol, 0.5% Triton X-100, 1 mM DTT, 1 mM EDTA, 1mM AEBSF, 20 µM leupeptin, 1 µM pepstatin, 1 mM PMSF). Fractions of 300 µL were collected at a flow rate of 0.5 mL/min. Fractions were precipitated by the addition of 4 volumes of cold acetone and stored overnight at -80°C. Proteins were pelleted by centrifugation at 21,000 g at 4°C for 20 min. Pellets were washed with cold acetone, air dried, resuspended in Laemmli buffer (10% glycerol, 2% SDS, 60 mM Tris-HCl pH 6.8, 0.02% bromophenol blue, 2.5% 2- mercaptoethanol) and were then subjected to SDS-PAGE and immunoblotting.

### Antibodies

The following antibodies were used for western blot and/or ChIP analyses: anti-HSP60 (sc13966, Santa Cruz Biotechnology), anti-H3K27ac (ab4729, Abcam), anti-H3K4me3 (ab8580, Abcam), anti-H3K79me2 (ab3594, Abcam), Anti-H3 total (ab1791, Abcam), anti- SET1A (clone-D3VPS, 61702S, Cell Signalling), anti-MLL1 (D2M7U, 14689S, Cell Signalling), anti-MLL2/KMT2B (D6X2E, 63735S, Cell Signalling), anti-Pol II (CTD4H8, 05- 623, Millipore), anti-FLAG M2 antibody (F3165, Sigma). Anti-ENL (ab245520, Abcam), anti-MLLT6 (PA5-30089, ThermoFisher), anti-MLLT10 (PA5-50907, ThermoFisher), anti- DOT1L (ab72454, Abcam) antibodies.

### qRT-PCR

mRNA was prepared with a Qiagen RNeasy kit and cDNA synthesis was performed with a SuperScript VILO kit (Life Technologies), per the manufacturers’ instructions. Quantitative PCR analysis was performed on an Applied Biosystems StepOnePlus System or Roche LightCycler 480 Real-Time PCR System with SYBR Green reagents. All samples were assayed in triplicate. Relative expression levels were determined with the ΔCt method and normalized to *GAPDH*. RT-qPCR primers are listed in Table 1.

### Cell proliferation and dose–response assays

For proliferation assays, cells were seeded at a consistent density prior to treatment in triplicate and treated with DMSO, VTP50469 or SGC0946 over the indicated time period. Drug was refreshed at least every three days. Cell number was calculated using the BD FACSverse (BD Biosciences), CytoFLEX (Beckman Coulter) or Muse cell counter.

### sgRNA competition assays

MLL-AF9 Cas9 cells were transduced with lentivirus expressing a gene specific sgRNA. The percentage of BFP-positive cells was measured between day 2 and day 12 after infection and normalised to the percentage of BFP at day 2.

### ChIP-sequencing

Chromatin immunoprecipitation was performed as described previously<u>50</u>. Briefly, for each ChIP, 20 million cells were crosslinked for 15 mins with 1% formaldehyde. Crosslinked material was sonicated to ∼200–1000 bp using the Covaris Ultrasonicator e220. Sonicated material was incubated overnight with each antibody, then incubated for 3 h with Protein A magnetic beads. Beads were washed with low and high salt wash buffers, LiCl buffer and TE, before being eluted and decrosslinked overnight. DNA was purified using Qiagen Minelute columns. All ChIP antibodies were used at ∼10ug per IP and are listed under the antibodies section. Sequencing libraries were prepared from eluted DNA using Rubicon ThruPLEX DNA-seq kit. Libraries were size selected between 200–500 bps and sequenced on the NextSeq500 using the 75 bp single-end chemistry. Chromatin from S2 cells was spiked-in to ChIP-seq experiments involving treatment with SGC0946 or knockout of DOT1L complex components. The S2 chromatin was added to chromatin from leukaemia cells prior to sonication at approximately 5-10% of the cell numbers used for the sample.

### ChIP-seq analysis

Reads were aligned to the human genome (GRCh37.73) with BWA-mem^51^. Duplicate reads and reads mapping to blacklist regions or the mitochondria were removed. Peak calling was performed with MACS2^52^ with default parameters. Genome-browser images of ChIP–seq data was generated by converting the bam files from BWA to TDF files with igvtools and viewing in IGV^53^. ChIP–seq coverage across selected genomic regions was calculated with BEDtools^54^. Average profile plots were generated using the NGSplot software ^75^.

### RNA-sequencing

RNA was extracted using the Qiagen RNeasy kit. RNA concentration was quantified using a Qubit Fluorometer (Thermo Fisher Scientific). Libraries were prepared using QuantSeq 3′ mRNA-seq Library Prep kit (Lexogen). Libraries were sequenced on the NextSeq500 using 75 bp single end chemistry.

### RNA-sequencing analysis

Bcl2fastq (Illumina) was used to perform sample demultiplexing and to convert BCL files generated from the sequencing instrument into Fastq files. Reads were aligned to the human genome (G1k V37) using HiSAT2^76^ and reads were assigned to genes using htseq-count^77^.

Differential expression was calculated using DESeq2^78^. Genes with a false discovery rate corrected for multiple testing using the method of Benjamini and Hochberg below 0.05 and a log fold change greater than 1 were considered significantly differentially expressed. Scatterplots depicting log fold change differences between conditions were generated in R using the ggplot2 package and custom R code.

### Quantification and statistical analysis

Statistical analysis was carried out using GraphPad Prism 9. Details of statistical analysis performed are provided in the figure legends. Data were reported as mean ± SD, SEM or independent replicates shown as individual data points, as indicated in the figure legends. Significance was defined as p < 0.05.

## Supplementary figure legends

**Extended Data Figure 1:**
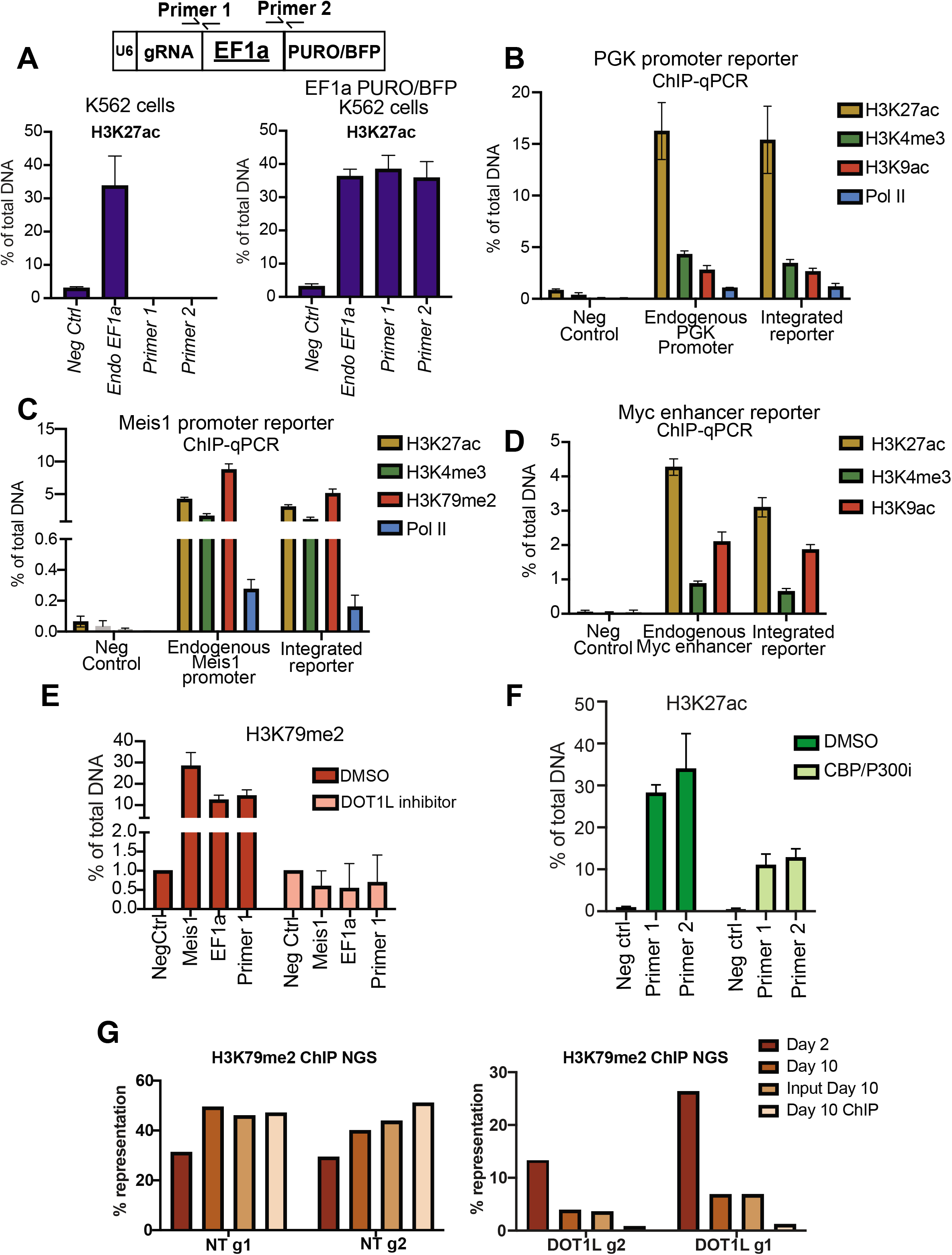
**A)** ChIP-qPCR analysis of H3K27ac ChIP performed in parental K562 cells or K562 cells infected with the CRISPR-ChIP plasmid containing an EF1a promoter. Primers for a negative control region (Neg Ctrl), endogenous EF1a or regions spanning the gRNA/EF1a (Primer 1), and EF1a/Puro (Primer 2) within the lentiviral CRISPR-ChIP vector were used, data represents mean +/- SD from two independent biological replicates. **B)** ChIP-qPCR analysis of H3K27ac, H3K4me3, H3K9ac and RNA-Pol II ChIP at the endogenous PGK promoter or integrated PGK promoter-Puro reporter, an intergenic region not enriched in active histone marks was used as a negative control (Neg control). Data represents mean +/- SD from two independent biological replicates. **C)** ChIP-qPCR analysis of H3K27ac, H3K4me3, H3K79me2 and RNA-Pol II ChIP at the endogenous Meis1 promoter or integrated Meis1 promoter (∼1.2Kb upstream of TSS)-Puro reporter, an intergenic region not enriched in active histone marks was used as a negative control (Neg control). Data represents mean +/- SD from two independent biological replicates. **D)** ChIP-qPCR analysis of H3K27ac, H3K4me3, and H3K9ac ChIP at the endogenous Myc enhancer or integrated Myc enhancer- mCMV-Puro sequence, an intergenic region not enriched in active histone marks was used as a negative control (Neg control). Data represents mean +/- SD from two independent biological replicates. **E)** ChIP-qPCR analysis of H3K79me2 in K562 cells infected with the CRISPR-ChIP plasmid and treated with DOT1L inhibitor (SGC0946) for 9 days. Primers for Negative control region (Neg Ctrl), Meis1, EF1a, and a region spanning the gRNA/EF1a were used, data represents mean +/- SD from two independent biological replicates. **F)** qPCR analysis of H3K27ac ChIP in K562 cells infected with the CRISPR-ChIP vector and treated with either DMSO or CBP/P300 inhibitor (A-485) for 24hrs. Primers for Negative control region (Neg Ctrl), Meis1, EF1a, and a region spanning the gRNA/EF1a were used, data represents mean +/- SD from two independent biological replicates. **G)** NGS representation of control guides (NT) at day 2, day 10, ChIP input day 10, and H3K79me2 ChIP (Left panel). NGS representation of DOT1L guides at day 2, day 10, ChIP input day 10, and H3K79me2 ChIP (Right panel).

**Extended Data Figure 2:**
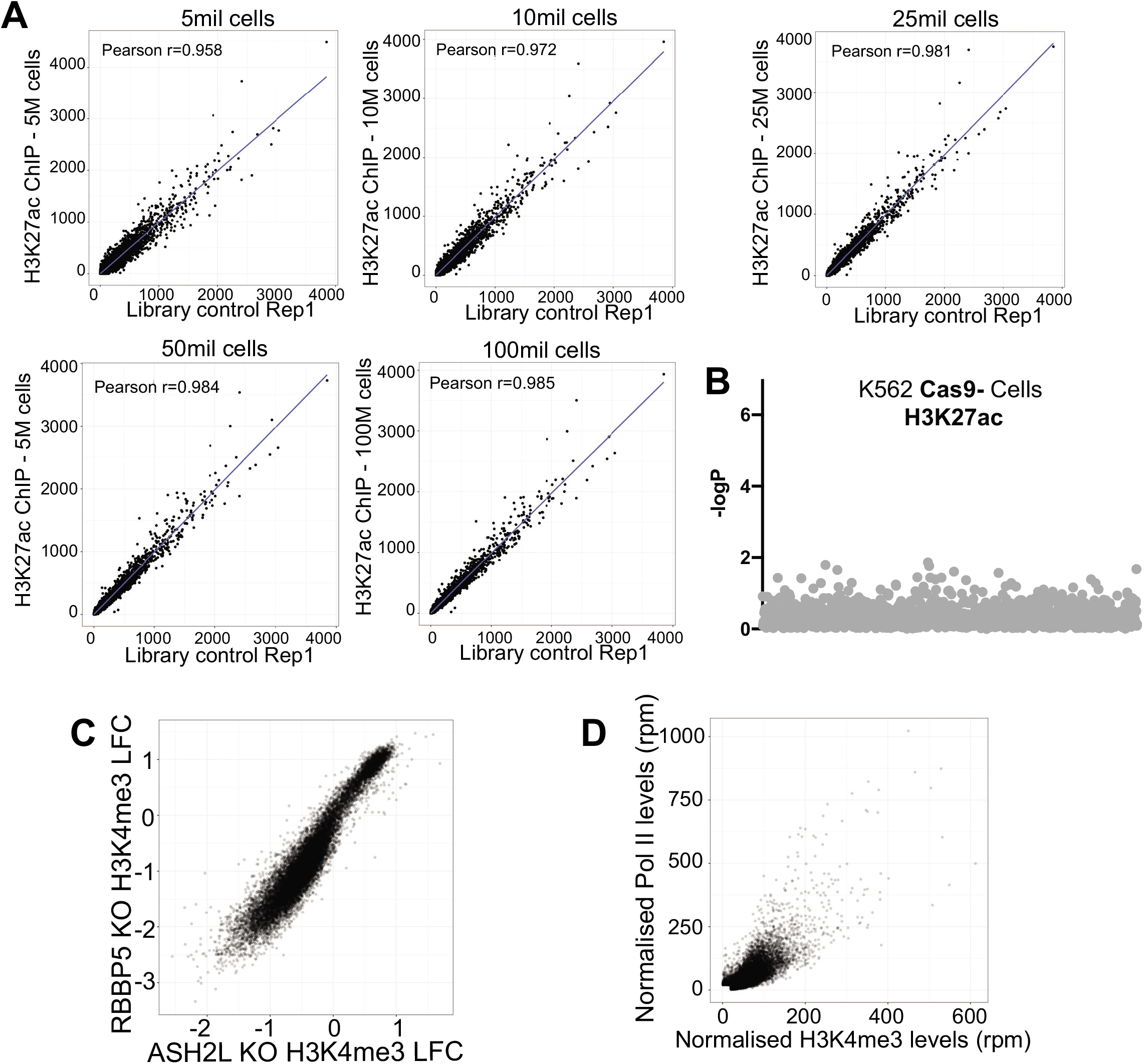
**A)** Correlation plot of library guide counts between H3K27ac ChIP and a library control sample taken from Cas9 negative K562 cells. H3K27ac ChIP was performed from different starting cell number, 5 million, 10 million, 25 million, 50 million and 100 million cells. **B)** Bubble plot of H3K27ac ChIP (50M cells) from Cas9 negative K562 cells analysed using MAGECK. **C)** Correlation plot of RBBP5 KO H3K4me3 LFC vs ASH2L KO H3K4me3 LFC. **D)** Correlation plot of normalised Pol II levels (rpm) vs normalised H3K4me3 levels (rpm).

**Extended Data Figure 3:**
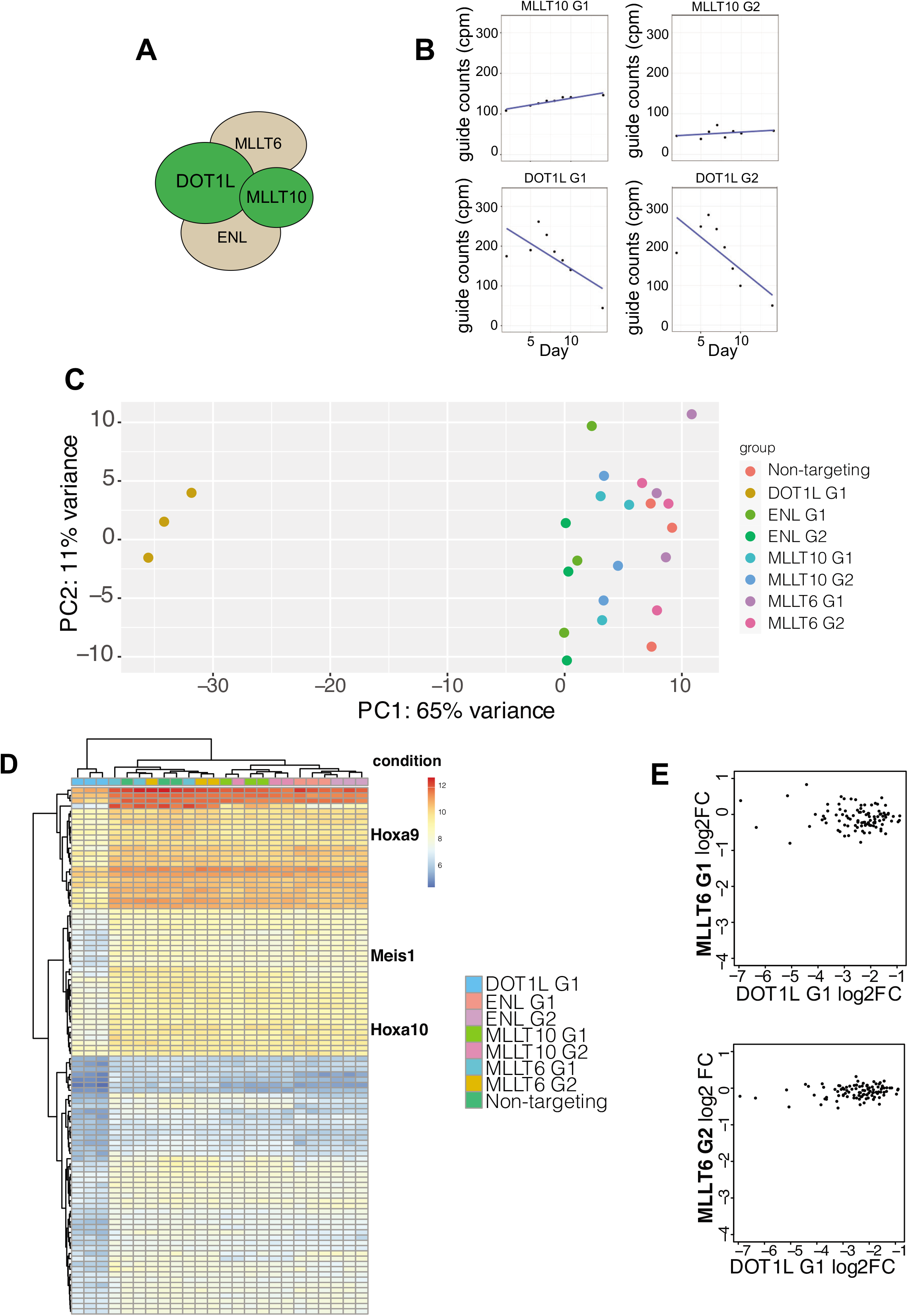
**A)** Schematic of DOTCOM complex (DOT1L, ENL/MLLT1, MLLT6/AF17, MLLT10). **B)** Guide counts over time from CRISPR dropout screen in MLL- AF9 cells for 2 MLLT10 and 2 DOT1L guides. **C)** PCA analysis of RNA-seq from MLLAF9 Cas9 cells transduced with control (non-targeting), DOT1L G1, two independent ENL guides, two independent MLLT10 guides and two independent MLLT6 guides. **D)** Heatmap of RNA-seq data described above, showing downregulated genes in the various knockouts as indicated. **E)** Correlation plot of RNA-seq LFC between two independent MLLT6 sgRNAs and DOT1L KO for the top 100 downregulated genes in the DOT1L KO cells.

**Extended Data Figure 4:**
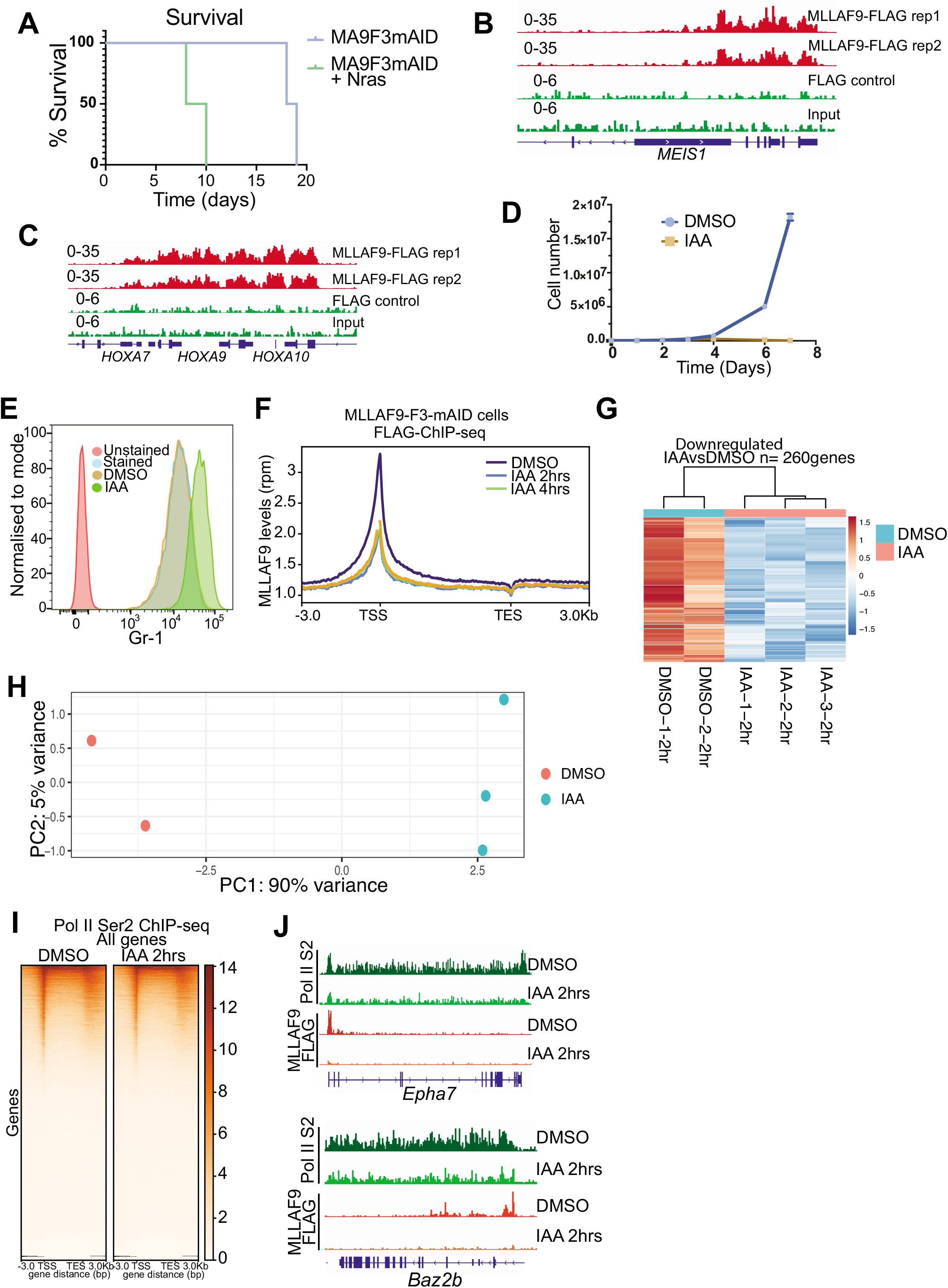
**A)** Kaplan-meir curve of mice transplanted with MLLAF9^FLAG-mAID^ cells or Nras + MLLAF9^FLAG-mAID^. **B)** Genome browser snapshot of MLLAF9 ChIP-seq replicates at the meis1 locus. FLAG ChIP from non-FLAG tagged MLLAF9 cells used as a negative control and input shown. **C)** Genome browser snapshot of MLLAF9 ChIP-seq replicates at the HOXA cluster. FLAG ChIP from non-FLAG tagged MLLAF9 cells used as a negative control and input shown. **D)** Proliferation assay of MLLAF9^FLAG-mAID^ Tir1 cells treated with DMSO or IAA (500uM) over 7 days. **E)** FACS analysis of cell surface Gr1 levels in unstained, stained, DMSO or IAA treated MLLAF9^FLAG-mAID^ Tir1 cells for 4 days. **F)** Profile plot of MLLAF9 FLAG ChIP-seq in MLLAF9^FLAG-mAID^ cells treated with DMSO, IAA (2hrs and 4hrs). **G)** Heatmap of 4su-seq (nascent RNA-seq) in MLLAF9^FLAG-mAID^ Tir1 cells treated with DMSO or IAA for 2hrs, showing the 260 downregulated genes. **H)** PCA plot of RNA-seq from MLLAF9^FLAG-mAID^ Tir1 cells treated with DMSO or IAA (2hrs). **I)** Heatmap of RNA Pol II ser2 ChIP-seq at all genes in MLLAF9^FLAG-mAID^ Tir1 cells treated with DMSO or IAA (2hrs). **J)** Genome browser snapshots of MLLAF9 targets, *Epha7* and *Baz2b* showing RNA Pol II- ser2 and MLLAF9 ChIP-seq tracks after treatment with either DMSO or IAA (2hrs).

**Extended Data Figure 5:**
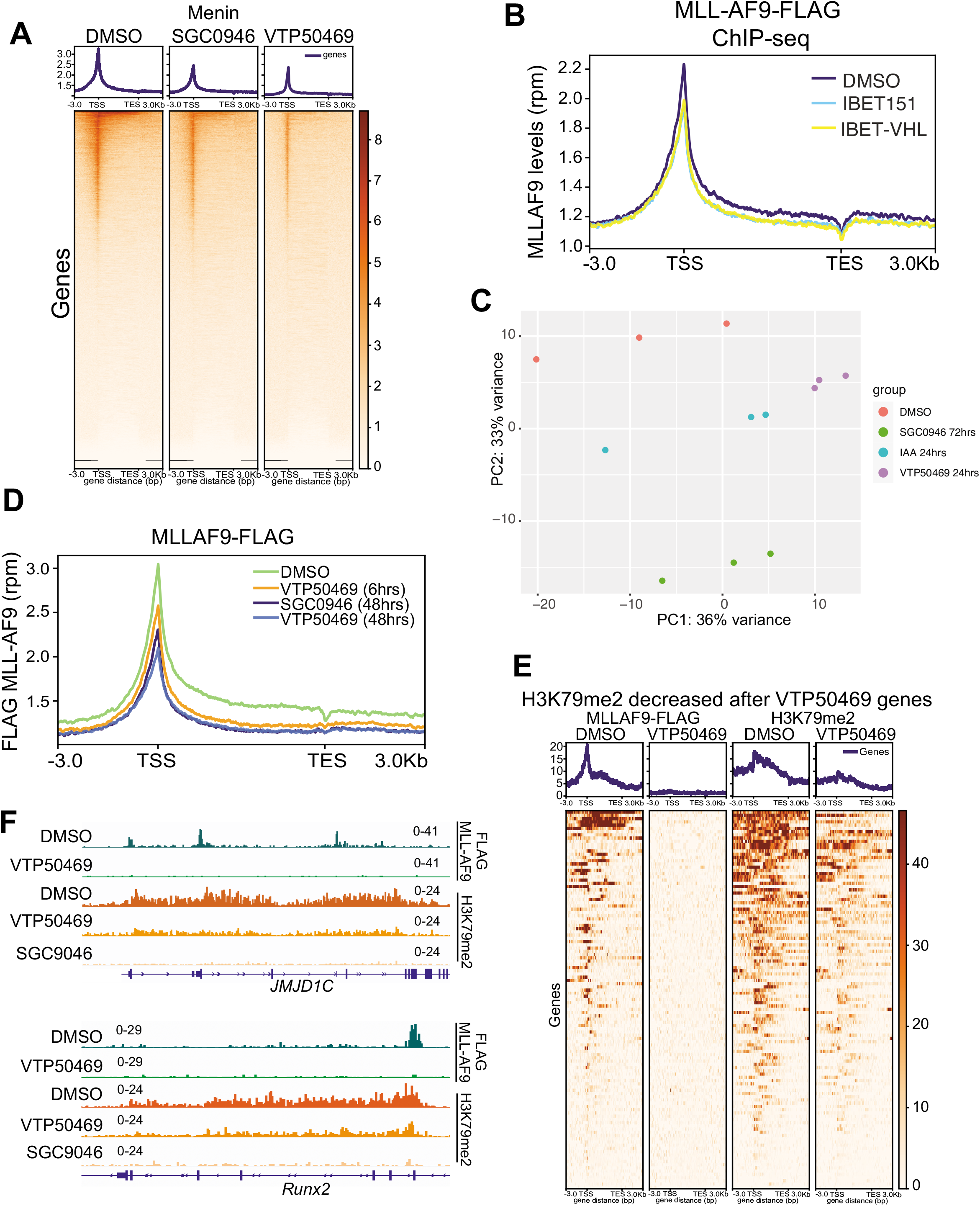
**A)** Heatmap of Menin ChIP-seq in MLLAF9 cells treated with DMSO, SGC0946 (72hrs) or VTP50469 (48hrs) at all genes. **B)** MLLAF9 FLAG ChIP-seq in MLLAF9 cells treated with DMSO, IBET151 or IBET-VHL for 8hrs at all genes. **C)** PCA analysis of RNA-seq data in MLLAF9 cells treated with DMSO, SGC0946 (72hrs), IAA (24hrs), VTP50469 (24hrs). **D)** Profile plot of MLLAF9 ChIP-seq in MLLAF9 cells treated with either DMSO, VTP50469 (6hrs), SGC0946 (48hrs) or VTP50469 (48hrs). **E)** Heatmap of MLLAF9 and H3K79me2 ChIP-seq data in MLLAF9 cells after treatment with VTIP50469 for 48hrs at genes that show decreased H3K79me2 after VTP50469 treatment. **F)** Genome browser snapshot of ChIP-seq tracks at canonical MLLAF9 target genes, *Jmjd1c* and *Runx2*, showing MLL-AF9 and H3K79me2 after treatment with VTP50469 or SGC0946 for 48hrs.

**Extended Data Figure 6:**
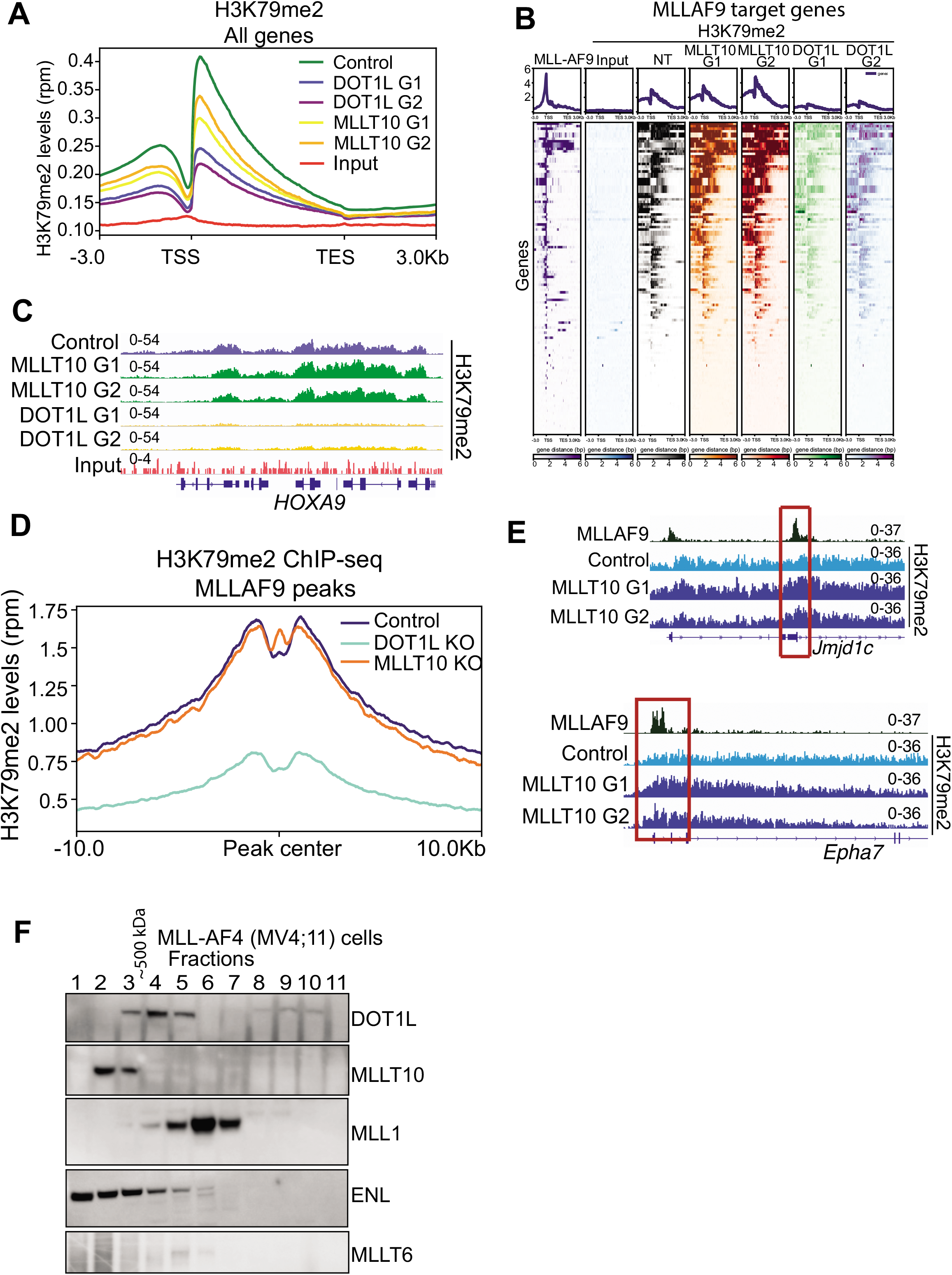
**A)** Average profile plot of H3K79me2 and input at all genes in MLL-AF9 cells infected with non-silencing guides (Control), 2 independent DOT1L guides and 2 independent MLLT10 guides. **B)** Heatmap of H3K79me2 at MLL-AF9 bound genes for the indicated samples. Genes are ranked from highest to lowest coverage for H3K79me2 in the control cells. **C)** Genome browser snapshot of a canonical MLLAF9 target, *Hoxa9*, from H3K79me2 ChIP-seq in MLLAF9 cells transduced with control sgRNA or two independent guides targeting MLLT10 or DOT1L. **D)** Profile plot of H3K79me2 distribution across MLLAF9 peaks +/- 10kb in MLLAF9^FLAG-mAID^ Cas9 cells transduced with control guides or guides targeting DOT1L or MLLT10. **E)** Genome browser snapshots of exemplar direct and strongly bound MLL-AF9 target genes that show increased H3K79me2 following MLLT10 KO in MLL-AF9 cells, *Jmjd1c and Epha7*. **F)** Immunoblot analysis of gel filtration fractions (superose 6 column) from nuclear extracts of MV4;11 cells (human MLL-AF4) using antibodies against DOT1L, MLLT10, MLL1, ENL and MLLT6.

**Extended Data Figure 7:**
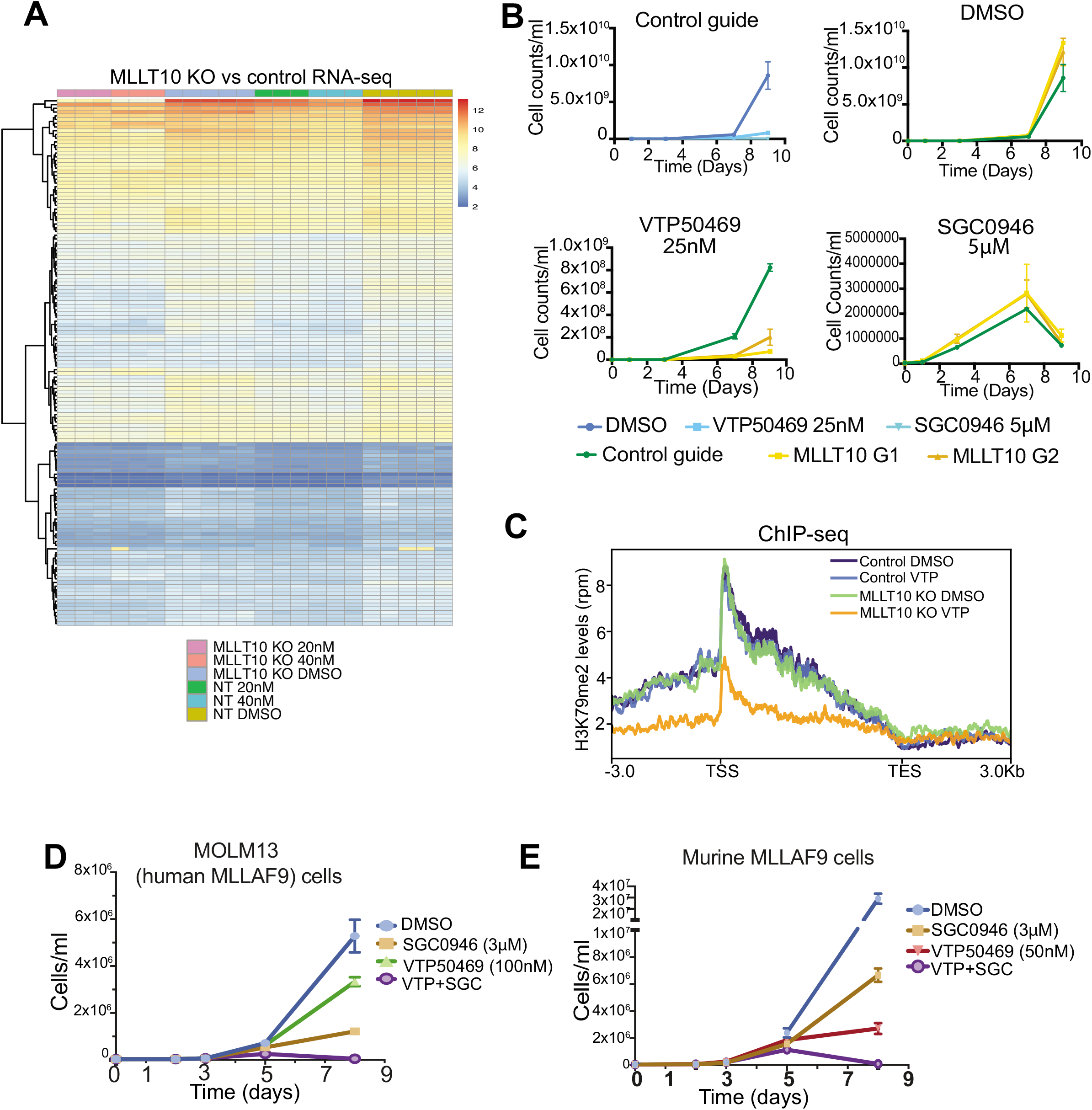
**A)** Heatmap of RNA-seq data from MLLAF9 Cas9 cells infected with control or MLLT10 guides and treated with either DMSO or VTP50469 with 20 or 40nM. **B)** Proliferation assays using MLL-AF9 Cas9 cells transduced with either a control non-targeting guide or two independent guides targeting MLLT10 (G1 and G2). Cells were treated with either DMSO, VTP50469 25nM or 50nM. Plots represent n=3 biological replicates. Bars represent mean +/- SD **C)** Profile plot of H3K79me2 ChIP-seq in MLLAF9 Cas9 cells infected with control or MLLT10 guides and treated with DMSO or VTP50469 (20nM) for 48hrs. Analysis performed across MLLAF9 target genes. **D)** Proliferation assay in MOLM13 cells treated with either DMSO, SGC0946 (3uM), VTP50469 (100nM) or combination. Mean +/- SD from n=3 biological replicates. **E)** Proliferation assay in MLLAF9^FLAG-mAID^ Cas9 cells treated with either DMSO, SGC0946 (3uM), VTP50469 (50nM) or combination. Cell counts shown at day 4 and day 6 and normalised to day 1 counts. Mean +/- SD from n=3 biological replicates. * p <0.05.

